# Mitochondria-Coupled Hitchhiking Drives Vimentin Intermediate Filament Transport

**DOI:** 10.64898/2026.11.02.763800

**Authors:** Bhuvanasundar Renganathan, Andrew Moore, Stephen A Adam, Jennifer Lippincott-Schwartz, Vladimir I. Gelfand

## Abstract

Vimentin intermediate filaments (VIFs) undergo active microtubule-dependent transport throughout the cell, with trajectories resembling those of motor-driven organelles, yet our recent work showed that only a small subset of VIFs lies close to microtubules. How VIFs engage the microtubule-based transport machinery, and whether additional transport routes contribute, remain unresolved. We hypothesized that VIFs move indirectly, by hitchhiking on motile organelles, and tested this using mitochondria. Live imaging showed mitochondria and adjacent VIFs moving together, and volume electron microscopy revealed VIFs closely following mitochondrial surfaces at sites distinct from microtubule contacts. Depleting the mitochondrial motor adaptor TRAK1 to inhibit anterograde mitochondrial transport shifted both mitochondria and VIFs toward the nucleus. Driving mitochondria toward the cell periphery with a tailless kinesin-1 in cells lacking endogenous kinesin-1, or toward the cell center with a minus-end-directed motor, redistributed a subset of VIFs in the same direction. Thus, VIF transport to a large extend is accomplished by hitchhiking on motile mitochondria.

## Introduction

Vimentin intermediate filaments (VIFs) are a core cytoskeletal component of mesenchymal and migratory cells.^1^ VIFs form a dense organized network that extends from the perinuclear region to the cell periphery VIFs interact physically and functionally with other cytoplasmic structures: they interpenetrate and interact with actin filaments ^2^ and microtubules,^3,4^ and they directly engage a wide range of membrane-bound organelles, including mitochondria, the endoplasmic reticulum, the Golgi apparatus, lipid droplets, and endosomes.^5–8^ Understanding how this filament network is itself organized and transported is therefore central to understanding how the cytoplasm as a whole is spatially structured.

Using single-particle tracking, we recently showed that VIFs are actively transported throughout the cell, with a constant fraction undergoing directed, microtubule-based motion irrespective of subcellular location, and with runs, pauses, and bidirectional trajectories resembling the transport of motor-driven organelles.^9^ Three-dimensional focused ion beam–scanning electron microscopy (FIB-SEM), however, showed that VIFs lie on average 233 nm from the nearest microtubule surface, with over 85% of microtubule length positioned more than 50 nm from any VIF.^9^

This creates a paradox. VIF transport and network organization depend on microtubules and their motors, kinesin-1 and cytoplasmic dynein,^10–12^ yet regions of close, extended VIF–microtubule contact are infrequent and spatially limited, making it unlikely that they alone support the full range of VIF motility. How, then, does VIF motility depend so strictly on microtubule motors without close microtubule proximity? Several non-exclusive mechanisms could explain this: brief, intermittent contacts between VIFs and microtubules may suffice for motor-driven transport; extended linker proteins such as plectin may bridge the gap between the two filament systems; or VIFs may move indirectly by hitchhiking, a mode of transport in which a cargo moves by associating with another cargo that has already recruited motors.^13,14^ Membrane-bound organelles are attractive candidates for such vehicles: many are transported along microtubules by kinesin-1 and cytoplasmic dynein, the same motors on which VIF motility depends,^15,16^ and VIFs are frequently found in close contact with organelles, in some cases forming cage-like structures around them.^17,18^ Hitchhiking on organelles could therefore explain how VIF motility depends on microtubule motors without close VIF–microtubule proximity.

To test the hitchhiking model, we used mitochondria as a representative organelle. The association of VIFs with mitochondria is well established, both physically and functionally. VIFs are linked to mitochondria through the cytolinker plectin isoform 1b^19^ and through a region of the vimentin N-terminal head domain, and they regulate mitochondrial motility and membrane potential.^5,6,20^ Moreover, mitochondrial transport along microtubules is driven by a well-characterized motor–adaptor complex, in which the outer-membrane GTPase MIRO1 recruits the adaptors TRAK1 and TRAK2, which engage kinesin-1 and dynein.^21–24^ Mitochondrial transport can therefore be selectively inhibited by depleting TRAK1 or redirected by recruiting motors to the mitochondrial outer membrane.^25^

Here, we combined live-cell imaging, super-resolution and volume electron microscopy, and targeted perturbations of mitochondrial transport to test whether VIFs hitchhike on mitochondria. We first asked whether mitochondria and VIFs move together and how they are organized relative to each other and to microtubules at the nanoscale. We then tested whether VIF distribution depends on mitochondrial transport, by depleting TRAK1, and whether mitochondrial movement is sufficient to reposition VIFs, by acutely biasing mitochondrial transport toward either microtubule end. Our results identify mitochondria as vehicles for a population of VIFs and support a hybrid model in which organelle hitchhiking acts alongside direct microtubule interactions to drive VIF motility.

## Results

### VIFs can move together with motile mitochondria

VIFs and mitochondria are both transported along microtubules, yet whether they move independently or are coupled during transport is unresolved. To test whether mitochondria and VIFs are co-transported in real time, we used spinning-disk confocal microscopy. We imaged mEmerald-vimentin knock-in RPE cells and labeled mitochondria with MitoTracker Deep Red. We observed instances of apparent transient co-motility of mitochondria and VIFs. In one example, a highly motile mitochondrion and adjacent VIFs changed position together over 36 s (Fig. 1 A, B and Movie 1). Thus, a mitochondrion and associated VIFs can move in a coordinated manner over tens of seconds. This observation motivated us to examine VIF–mitochondria interactions at higher resolution.

**Figure 1.**
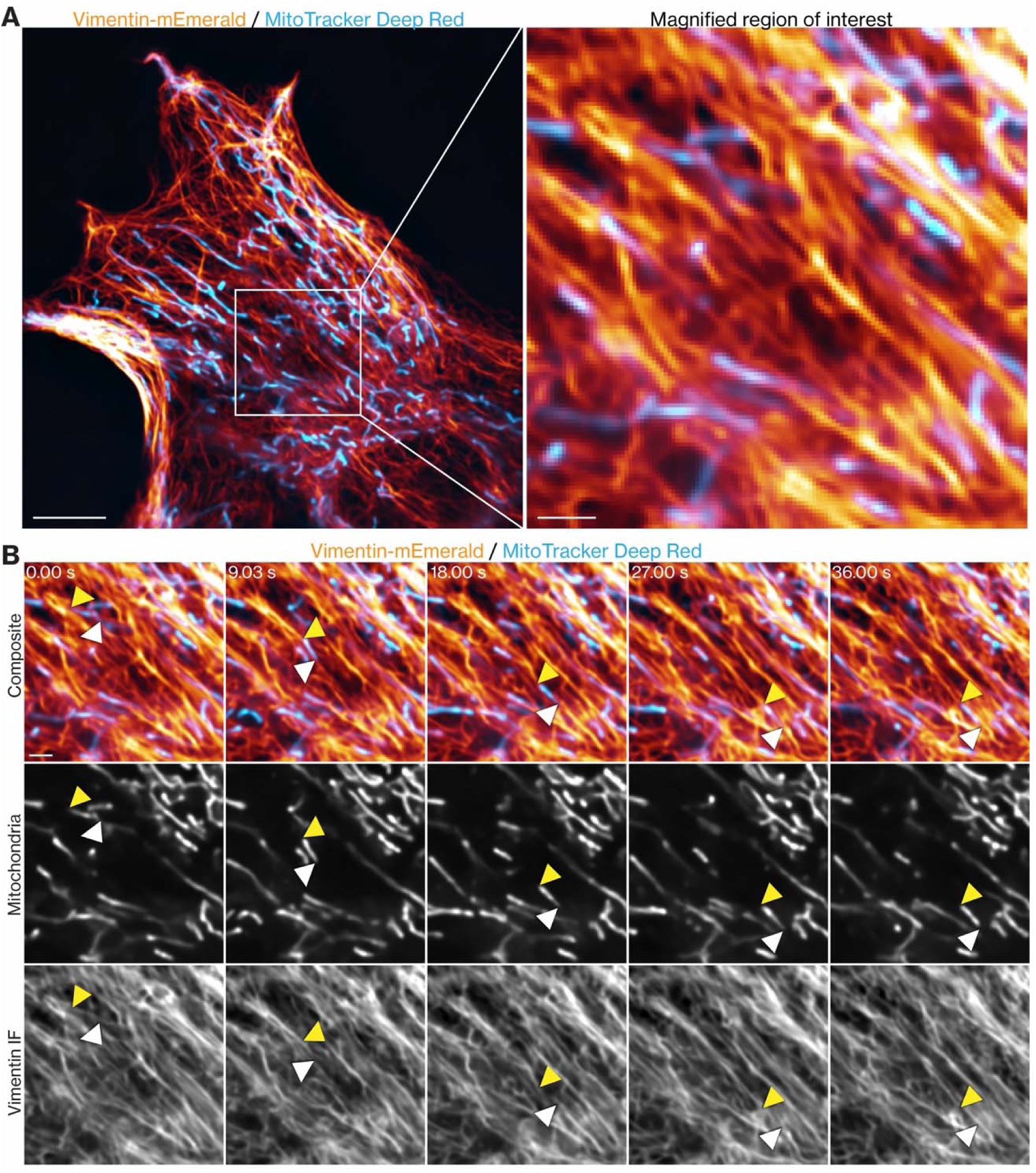
VIFs can move together with a motile mitochondrion. (A) Spinning-disk confocal image of a live mEmerald-vimentin knock-in RPE cell (orange) labeled with MitoTracker Deep Red (blue). The boxed region is magnified at right. (B) Time-lapse sequence of the boxed region over 36 s, showing co-motility of a mitochondrion with adjacent VIFs. Two-color merges are shown above grayscale single-channel panels. White and yellow arrowheads mark the leading and trailing ends of the mitochondrion, respectively. See also Movie 1. Scale bars: 10 µm (A, left); 2 µm (A, right; B).

### Mitochondria are located within intersecting VIF and microtubule networks

We next asked whether VIFs associate with mitochondria independently of microtubules, or only where both structures share a common microtubule track to which each could be independently linked. To address this, we performed STED microscopy on fixed COS-7 cells immunolabeled for vimentin, α-tubulin, and the mitochondrial outer membrane protein TOM20; vimentin and α-tubulin were imaged by STED, and TOM20 by confocal microscopy. The multicolor images revealed mitochondria positioned within an extensive, crisscrossing cytoskeletal network marked by frequent sites of overlap between mitochondria, microtubules, and VIFs (Fig. 2). We also observed sites where VIFs crossed mitochondria without a detectable microtubule in the imaged plane (Fig. 2, arrows). However, the precise nanoscale organization of these sites could not be resolved, even with the enhanced resolution offered by STED microscopy. We therefore turned to volume electron microscopy.

**Figure 2.**
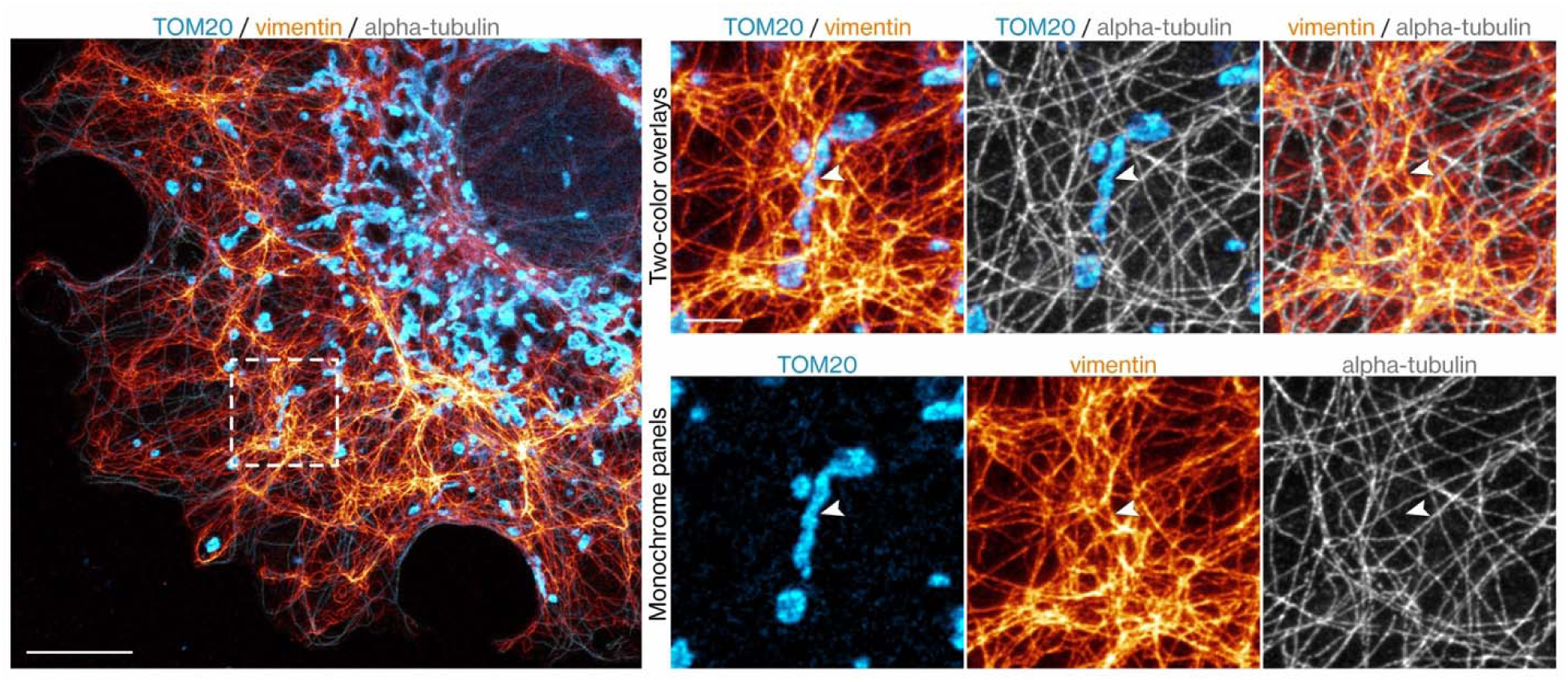
Mitochondria are located within intersecting VIF and microtubule networks. STED microscopy of a fixed COS-7 cell immunostained for TOM20 (mitochondria, light blue), vimentin (orange), and α-tubulin (microtubules, gray). The dotted boxed region is enlarged at right, with pairwise merged channels in the upper row and single channels below. Arrows indicate a site where VIFs cross a mitochondrion without a detectable microtubule in the imaged plane. Vimentin and α-tubulin were imaged by STED and TOM20 by confocal microscopy. Scale bars: 10 µm (whole cell); 2 µm (insets).

### A subset of VIFs closely traces mitochondrial surfaces in volume EM reconstructions

To comprehensively reconstruct VIF-mitochondria contacts at sufficient 3D spatial resolution to infer structural relationships, we turned to volume EM. Specifically, we reanalyzed a high-resolution FIB-SEM volume in the perinuclear region of a COS-7 cell from which we had previously reconstructed individual cytoskeletal fibers: 572 VIFs and 39 microtubules.^6^ Here, we traced the surface of 15 mitochondrial profiles present in the volume. From the 3D segmentation masks, we noted that mitochondria lay nestled within the VIF meshwork, with several conspicuous examples of individual filaments curving along portions of the mitochondrial surface (Fig. 3, A–C; Fig. S1, A-C; Movie 2).

**Figure 3.**
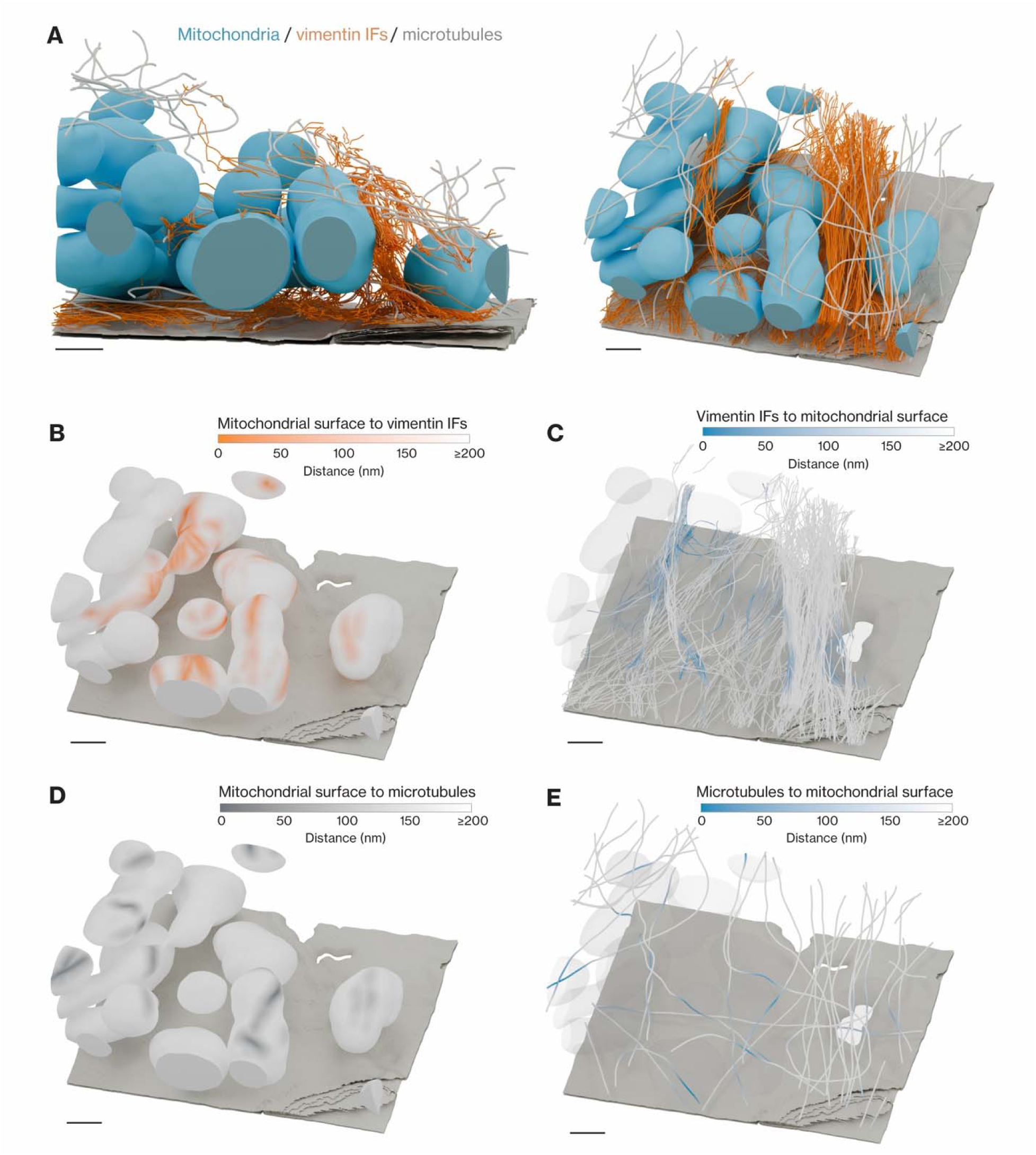
A subset of VIFs closely traces mitochondrial surfaces in volume EM reconstructions. (A) Two orthographic views of mitochondria (light blue), VIFs (orange), and microtubules (light gray) reconstructed from a FIB-SEM volume of a COS-7 cell. The segmented basal plasma membrane (gray) provides anatomical context. (B and C) Reciprocal distance maps showing mitochondria-to-VIF distance displayed on mitochondria (B) and VIF-to-mitochondria distance displayed on VIFs (C). (D and E) Corresponding maps for microtubules, showing mitochondria-to-microtubule distance displayed on mitochondria (D) and microtubule-to-mitochondria distance displayed on microtubules (E). Colors fade to white at distances of 200 nm or greater. All maps derive from one FIB-SEM volume. See also Movie 2. Scale bars, 500 nm.

To describe this proximity, we measured distances between the native mitochondrial segmentation and filament centerlines modeled as tubes of 6 nm radius for vimentin and 12 nm for microtubules. In this reconstructed cell volume, 4.61% of the analyzed mitochondrial surface lay within 50 nm of a vimentin filament, compared with 1.15% within the same distance of a microtubule (Fig. S1E). Measured along the filaments, 3.87% of the analyzed vimentin length and 4.98% of the microtubule length lay 0–50 nm from the mitochondrial surface (Fig. S1F).

### Reciprocal maps distinguish local VIF and microtubule proximity

We also mapped mitochondrial proximity to microtubules in both directions, using the same views and distance range as for VIFs (Fig. 3, D and E; Fig. S1D, Movie 3). These complementary maps indicated where proximal microtubules were positioned along the mitochondrial surface and where along each microtubule the proximity to mitochondria was greatest. These maps reveal sites of close association between mitochondria and microtubules that do not universally correspond to sites of VIF–mitochondrial apposition.

### Inhibition of mitochondrial transport affects VIF distribution

TRAK1 is a mitochondrial motor adaptor that is recruited to the outer mitochondrial membrane by MIRO1, links mitochondria to kinesin-1, and is required for kinesin-1-dependent anterograde mitochondrial transport. TRAK1 loss inhibits mitochondrial transport^21^ and, in non-neuronal cells, shifts mitochondria from a kinesin-1-driven peripheral distribution toward perinuclear accumulation.^23^ To test whether disrupting this pathway also shifts VIFs toward the perinuclear region, we depleted TRAK1 in RPE cells by transient shRNA-mediated knockdown and compared mitochondrial and VIF distributions with those in control RPE cells by immunofluorescence (Fig. 4; Fig. S2).

**Figure 4.**
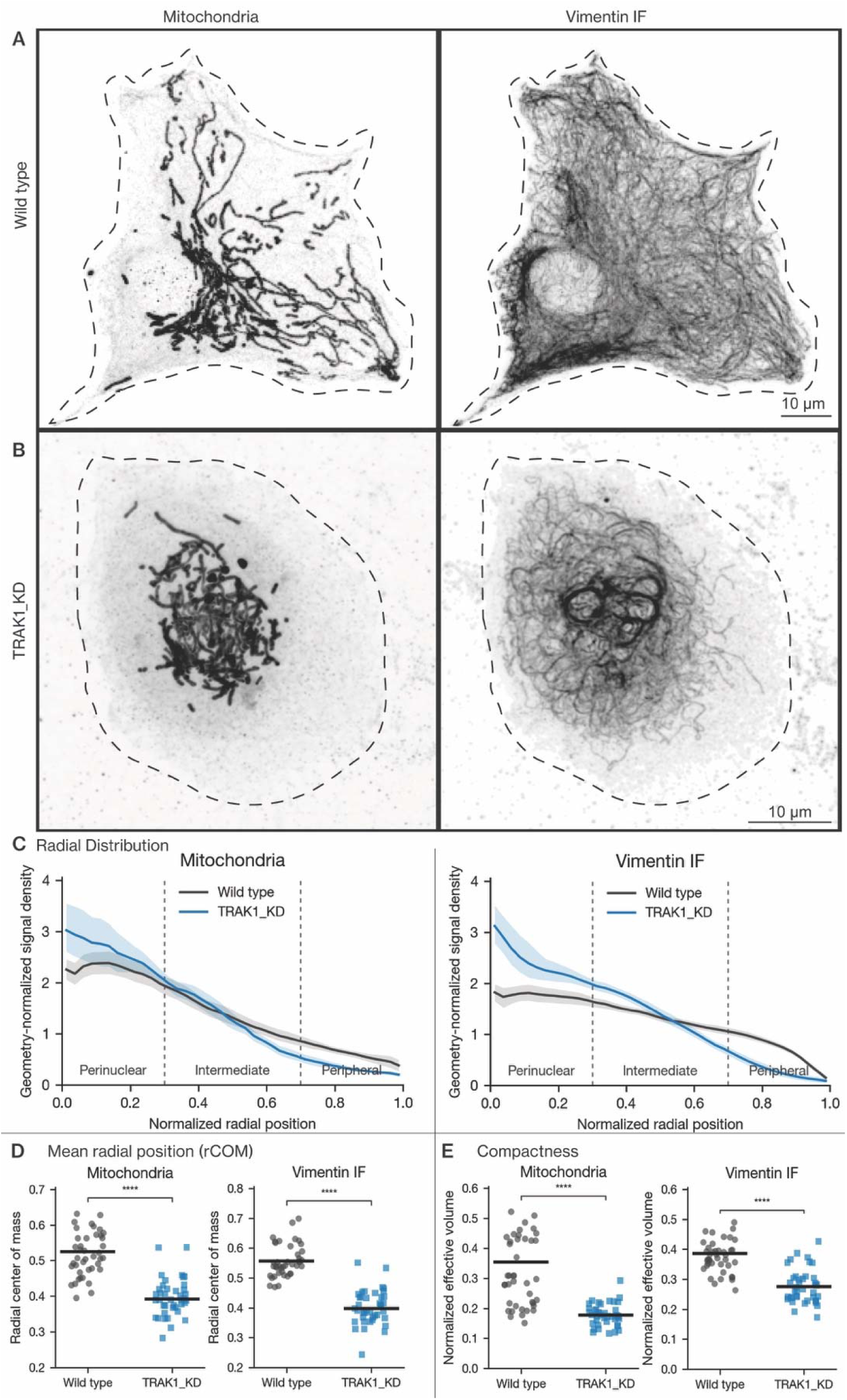
TRAK1 depletion in RPE cells shifts mitochondria and VIFs toward the perinuclear region. (A and B) Representative inverted-contrast immunofluorescence images of mitochondria (left) and VIFs (right) in wild-type (A) and TRAK1 KD (B) RPE cells. Dashed lines outline the cell boundary. (C) Geometry-normalized radial profiles of mitochondria (left) and VIFs (right) as a function of normalized radial position (0, nuclear boundary; 1, cell edge) in wild-type (black) and TRAK1 KD (blue) cells. (D) Mean radial position (rCOM) of mitochondria (left) and VIFs (right). (E) Compactness (normalized effective intensity volume) of mitochondria (left) and VIFs (right). Each point is one cell; bars show the mean. Regional signal fractions are shown in Fig. S2. Scale bar, 10 µm. N = 41 wild-type and 40 TRAK1 KD cells from three independent experiments.

TRAK1 KD transformed the extended, branched mitochondrial network in control RPE cells into a dense perinuclear cluster (Fig. 4, A and B). Geometry-normalized 3D radial intensity profiling, which compares the share of fluorescence in each radial shell with that shell’s share of cytoplasmic volume (see Materials and methods), confirmed a pronounced redistribution toward the perinuclear region and away from the cell periphery (Fig. 4C, left). VIFs underwent a parallel reorganization: the extended network in control cells collapsed into a dense perinuclear cage after TRAK1 KD, with a corresponding inward shift in the radial profile (Fig. 4C, right).

To quantify these changes, we summarized each distribution by its mean radial position (the intensity-weighted average position between the nuclear boundary, 0, and the cell edge, 1), its regional signal fractions (perinuclear, 0–0.3; intermediate, 0.3–0.7; peripheral, 0.7–1.0), and its compactness (normalized effective intensity volume, in which lower values indicate more concentrated signal regardless of position). Consistent with the profiles, mitochondrial mean radial position decreased significantly from 0.52 in control cells to 0.40 after TRAK1 KD, and the peripheral mitochondrial fraction fell from 30.9% to 15% (Fig. 4D, left; Fig. S2A). VIFs showed a similar inward redistribution, with mean radial position decreasing significantly from 0.56 to 0.40 and the peripheral VIF fraction falling from 36.0% to 13.5% (Fig. 4D, right; Fig. S2B). In addition to this change in radial position, both networks became more compact: normalized effective intensity volume decreased significantly from 0.32 to 0.18 for mitochondria and from 0.37 to 0.28 for VIFs (Fig. 4E). Together, these measurements show that TRAK1 depletion causes a coordinated inward redistribution and compaction of both the mitochondrial and VIF networks.

To determine whether this coordinated redistribution was a general consequence of TRAK1 loss rather than an RPE-specific response, we generated stable TRAK1 KD BJ05 fibroblasts (Fig. 5; Fig. S3). As in RPE cells, the mitochondrial network collapsed into a dense perinuclear cluster (Fig. 5, A and B, middle), and its mean radial position decreased significantly from 0.51 to 0.44 (Fig. 5, C and D, left; Fig. S3A). VIFs again collapsed into a perinuclear cage (Fig. 5, A and B, right), with a larger decrease in mean radial position, from 0.57 to 0.41 (Fig. 5, C and D, right; Fig. S3B). Both networks also became significantly more compact (Fig. 5E). Microtubules continued to extend from the perinuclear region to the cell periphery (Fig. 5, A and B, left), indicating that the mitochondrial and VIF redistribution was not secondary to loss of microtubule tracks.

**Figure 5.**
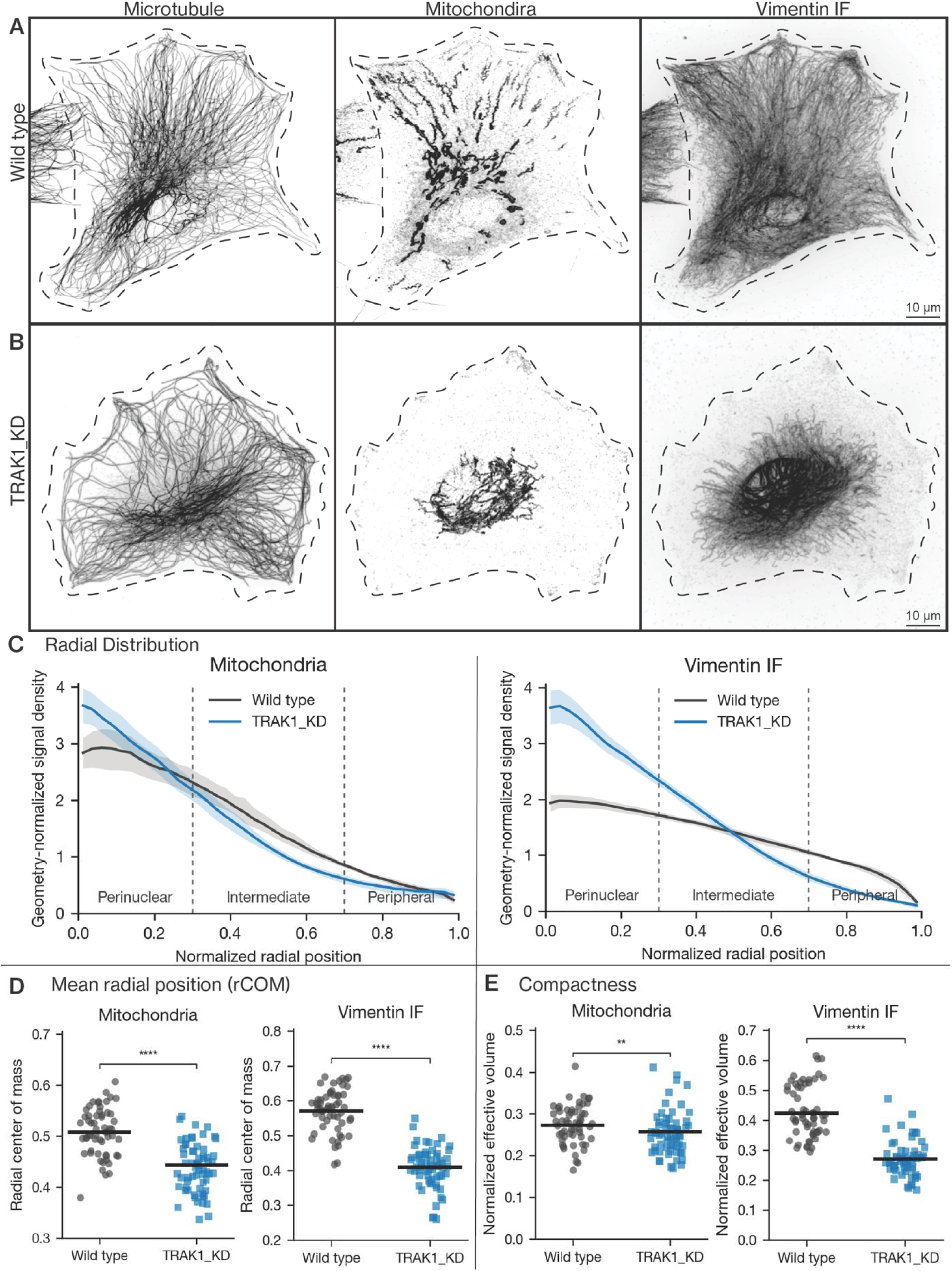
Stable TRAK1 depletion in BJ05 fibroblasts reproduces the perinuclear shift of mitochondria and VIFs. (A and B) Representative inverted-contrast immunofluorescence images of microtubules (left), mitochondria (middle), and VIFs (right) in wild-type (A) and TRAK1 KD (B) BJ05 cells. Dashed lines outline the cell boundary. (C) Geometry-normalized radial profiles of mitochondria (left) and VIFs (right) in wild-type (black) and TRAK1 KD (blue) cells. (D) Mean radial position (rCOM) of mitochondria (left) and VIFs (right). (E) Compactness (normalized effective intensity volume) of mitochondria (left) and VIFs (right). Regional signal fractions are shown in Fig. S3. Scale bar, 10 µm. N = 58 wild-type and 63 TRAK1 KD cells from three independent experiments.

As an independent test at the level of the motor, we examined RPE cells lacking kinesin-1 (KIF5B knockout [KO]^12^) (Figs. S4 and S5). Kinesin-1 loss reproduced the coordinated perinuclear shift: mean radial position decreased significantly from 0.52 to 0.45 for mitochondria and from 0.54 to 0.41 for VIFs (Fig. S4, C and D), whereas the radial distribution of microtubules did not change significantly (Fig. S4, A and B, left), indicating that the microtubule tracks themselves were not disrupted. Both perturbations therefore shifted mitochondria and VIFs toward the perinuclear region. The relative magnitudes, however, differed between the two networks. Mitochondria were displaced more strongly after TRAK1 KD than after kinesin-1 KO (mean radial position 0.52 to 0.39, versus 0.52 to 0.45), whereas VIFs were displaced to a similar extent in both (0.56 to 0.39, versus 0.54 to 0.41; Fig. 4D; Fig. S4D). This descriptive comparison suggests that the magnitude of VIF redistribution does not simply scale with the magnitude of mitochondrial redistribution.

Together, these results show that inhibiting kinesin-1/TRAK1-dependent mitochondrial transport shifts both mitochondria and VIFs toward the perinuclear region. The same coupled shift in RPE and BJ05 cells indicates that it is not cell-type specific, and in BJ05 and kinesin-1 KO cells it occurred without loss of the microtubule network. These findings extend prior evidence that vimentin and mitochondria are physically and functionally coupled^6,18,20,26^ by showing that kinesin-1/TRAK1-dependent mitochondrial transport contributes to the peripheral distribution of VIFs.

### Biasing mitochondrial transport repositions VIFs

#### Recruiting kinesin-1 to mitochondria shifts VIFs toward the cell periphery

Loss of TRAK1 or kinesin-1 shifted mitochondria and VIFs toward the perinuclear region together (Figs. 4 and 5; Figs. S2–S5), and live imaging showed that the two structures can move together (Fig. 1 and movie 1). These experiments did not establish, however, whether moving mitochondria is sufficient to reposition VIFs. To test this, we used chemically induced dimerization to recruit a constitutively active kinesin-1 motor directly to mitochondria.^25^ In kinesin-1 KO RPE cells, which lack endogenous kinesin-1, we co-expressed a kinesin-1 heavy-chain fragment comprising only the motor domain (residues 1–560; hereafter, truncated kinesin-1) fused to FRB, together with FKBP targeted to the outer mitochondrial membrane (OMM–FKBP) (Fig. 6A). Truncated kinesin-1 lacks the tail region required for kinesin-1-dependent IF transport.^12^ Addition of the rapalog A/C Heterodimerizer (hereafter, dimerizer) for 2 h therefore directed kinesin-1 activity specifically to mitochondria, bypassing the endogenous TRAK/MIRO adaptor complex.

**Figure 6.**
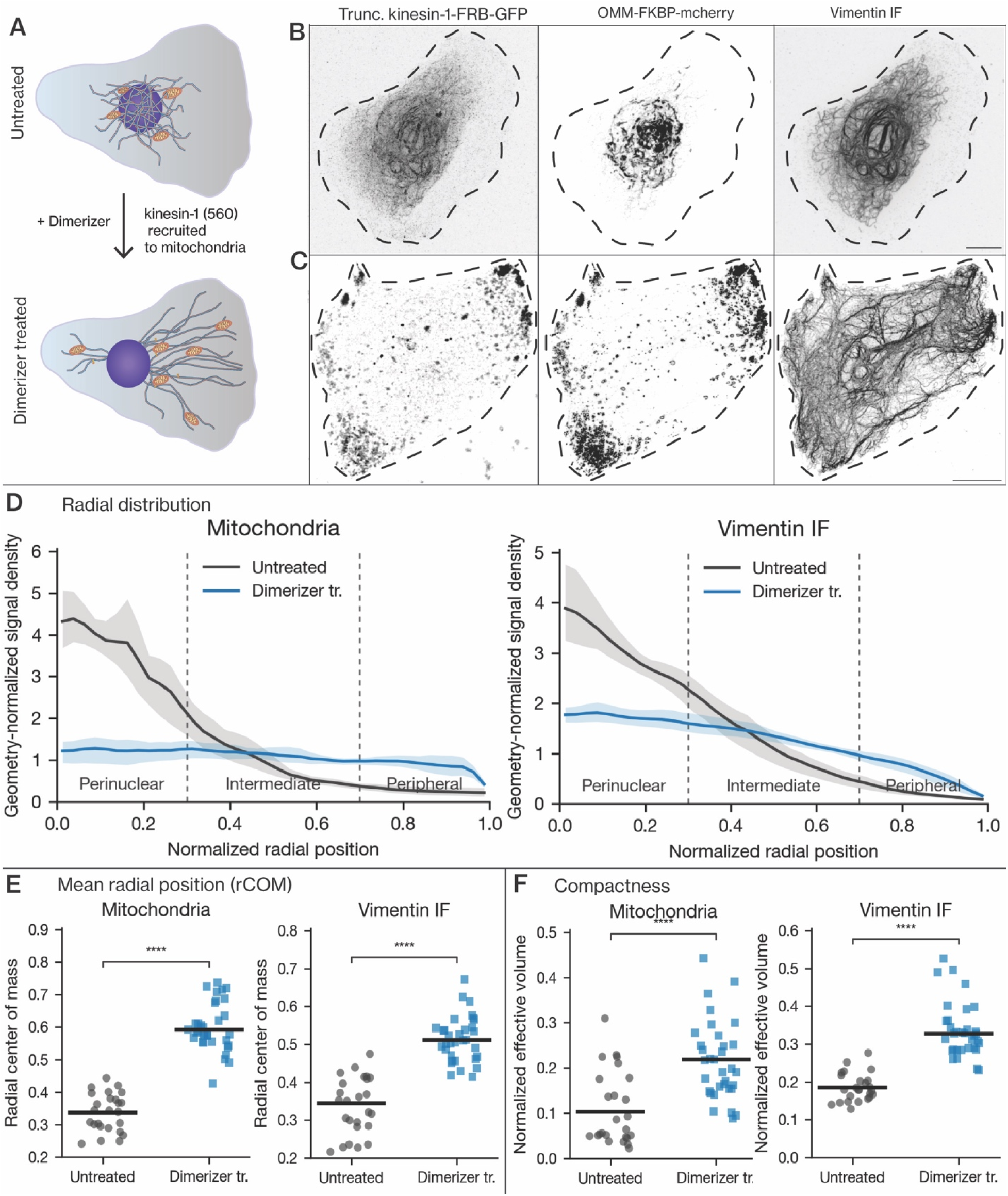
Recruiting truncated kinesin-1 to mitochondria shifts VIFs toward the cell periphery. (A) Schematic of the dimerization strategy. Truncated kinesin-1 (KIF5B residues 1–560, lacking the cargo-binding tail) fused to FRB is recruited by the dimerizer to FKBP targeted to the outer mitochondrial membrane (OMM–FKBP), directing kinesin-1 activity to mitochondria independently of the endogenous TRAK/MIRO adaptor complex. (B and C) Representative inverted-contrast images of truncated kinesin-1–FRB–GFP (left), OMM–FKBP–mCherry-labeled mitochondria (middle), and VIFs (right) in kinesin-1 KO RPE cells expressing both constructs, without (B) or with (C) dimerizer for 2 h. Dashed lines outline the cell boundary. (D) Geometry-normalized radial profiles of mitochondria (left) and VIFs (right) in untreated (black) and dimerizer-treated (blue) cells. (E) Mean radial position (rCOM) of mitochondria (left) and VIFs (right). (F) Compactness (normalized effective intensity volume) of mitochondria (left) and VIFs (right). Each point is one cell; bars show the mean. Regional signal fractions are shown in Fig. S6. Scale bar, 10 µm. N = 25 untreated and 33 dimerizer-treated cells from three independent experiments.

Without dimerizer, truncated kinesin-1 remained diffuse, and both mitochondria and VIFs retained the perinuclear distribution characteristic of kinesin-1 KO cells (Fig. 6B). Dimerizer recruited truncated kinesin-1 to mitochondria and redistributed mitochondria toward the cell periphery (Fig. 6C, middle), shifting their radial profile away from the perinuclear region (Fig. 6D, left); mitochondrial mean radial position increased significantly from 0.34 to 0.60 (Fig. 6E, left; Fig. S6A). VIFs redistributed in parallel, extending from the perinuclear region into the peripheral cytoplasm (Fig. 6C, right), with a corresponding outward shift in their radial profile (Fig. 6D, right) and a significant increase in mean radial position from 0.34 to 0.51 (Fig. 6E, right; Fig. S6B). Both networks also became significantly less compact (Fig. 6F and S7), consistent with their spreading through the cytoplasm.

Because truncated kinesin-1 was recruited to the mitochondrial outer membrane, lacks the cargo-binding tail and therefore cannot bind cargoes on its own, and was expressed in cells lacking endogenous kinesin-1, the VIF redistribution is unlikely to reflect direct engagement of VIFs by kinesin-1. Instead, these results indicate that redirecting mitochondrial transport is sufficient to shift VIFs toward the cell periphery.

#### Recruiting a minus-end-directed motor to mitochondria shifts VIFs toward the nucleus

We next asked whether biasing mitochondrial transport in the opposite direction would shift VIFs toward the nucleus. To drive retrograde transport, we used kinesin-14 VIb from the moss *Physcomitrella patens* (hereafter, kinesin-14) minus-end-directed motor, which has previously been used in mammalian cells to move organelles toward microtubule minus ends.^27^ In wild-type RPE cells, in which endogenous kinesin-1 is intact, we recruited FRB-tagged kinesin-14 VIb to OMM–FKBP with the dimerizer (Fig. 7A and S9), thereby biasing mitochondrial transport toward microtubule minus ends near the nucleus.

**Figure 7.**
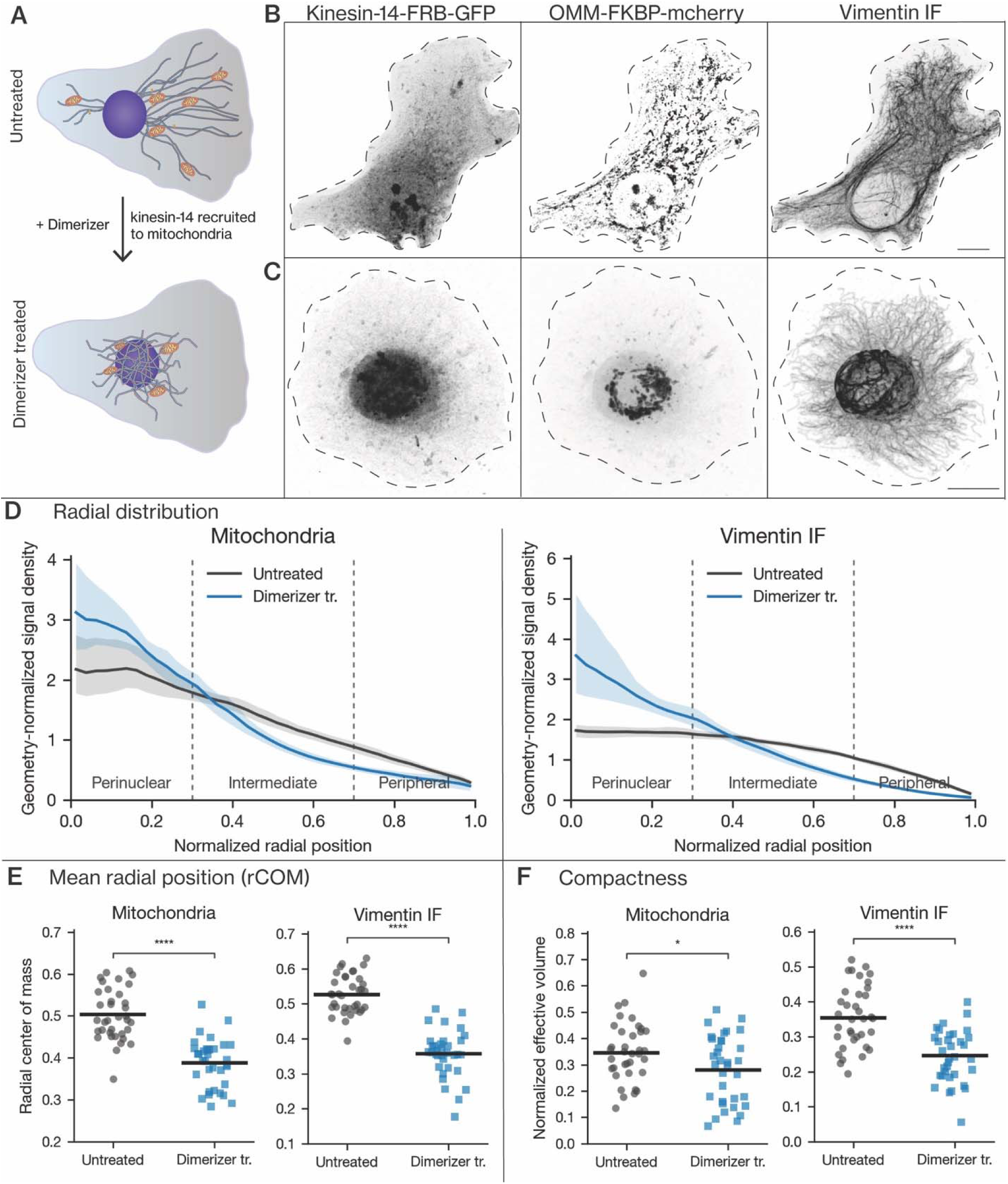
Recruiting kinesin-14 VIb to mitochondria shifts a substantial fraction of VIFs toward the nucleus. (A) Schematic of the dimerization strategy. The minus-end-directed motor kinesin-14 VIb fused to FRB is recruited by the dimerizer to OMM–FKBP in wild-type RPE cells, biasing mitochondrial transport toward microtubule minus ends. (B and C) Representative inverted-contrast images of kinesin-14 VIb–FRB–GFP (left), OMM–FKBP–mCherry-labeled mitochondria (middle), and VIFs (right) in cells expressing both constructs, without (B) or with (C) dimerizer. Dashed lines outline the cell boundary. (D) Geometry-normalized radial profiles of mitochondria (left) and VIFs (right) in untreated (black) and dimerizer-treated (blue) cells. (E) Mean radial position (rCOM) of mitochondria (left) and VIFs (right). (F) Compactness (normalized effective intensity volume) of mitochondria (left) and VIFs (right). Each point is one cell; bars show the mean. Regional signal fractions are shown in Fig. S8. Scale bar, 10 µm. N = 36 untreated and 34 dimerizer-treated cells from three independent experiments.

Without dimerizer, kinesin-14 was diffuse, mitochondria were dispersed throughout the cytoplasm, and VIFs extended from the perinuclear region to the cell periphery (Fig. 7B). Dimerizer recruited kinesin-14 to mitochondria and produced a tight perinuclear mitochondrial cluster (Fig. 7C, middle), with increased perinuclear and reduced peripheral signal in the radial profile (Fig. 7D, left). Mitochondrial mean radial position decreased significantly from 0.50 to 0.39, and the peripheral mitochondrial fraction fell from 28.4% to 16.6% (Fig. 7E, left; Fig. S8A). Retrograde recruitment thus reproduced the perinuclear mitochondrial distribution seen after loss of TRAK1 or kinesin-1 (Figs. 4 and 5; Figs. S2–S5), the reverse of the anterograde manipulation (Fig. 6).

VIFs also formed a dense perinuclear aggregate after dimerization (Fig. 7C, right), and their mean radial position decreased significantly from 0.53 to 0.36 (Fig. 7E, right; Fig. S8B). The VIF radial profile, however, did not fully match the mitochondrial profile: peripheral VIF signal was depleted less than peripheral mitochondrial signal (Fig. 7D; Fig. S7, A and B), a difference that mean radial position alone does not capture, because the dense perinuclear aggregate lowers the mean even while a peripheral VIF population persists. Compactness also responded differently: mitochondrial compactness changed only modestly (normalized effective intensity volume, 0.35 to 0.28; P < 0.05), whereas VIFs became markedly more compact (0.36 to 0.24; P < 0.0001) (Fig. 7F and S9).

Thus, retrograde mitochondrial transport shifted a substantial fraction of VIFs toward the nucleus, while a peripheral VIF population remained. This partial divergence indicates that not all VIFs follow repositioned mitochondria, consistent with a subset of VIFs being coupled to mitochondria while others are positioned by additional mechanisms.

## Discussion

Our previous single-particle tracking showed that individual VIFs undergo microtubule-dependent directed motion yet rarely make close, extended contact with microtubules.^9^ Here, we tested whether motile organelles provide an indirect route for this motility, using mitochondria as a representative cargo. VIFs and mitochondria moved together in living cells, and VIFs closely followed mitochondrial surfaces at sites distinct from mitochondria–microtubule contacts. Inhibiting mitochondrial transport shifted VIFs toward the nucleus together with mitochondria, and redirecting mitochondria toward either microtubule end shifted VIFs in the same direction. Together, these findings support a model in which a subset of VIFs moves along microtubules indirectly, by associating with motile mitochondria.

The dependence of VIF transport and organization on microtubules and their motors has been recognized for decades. Microtubule depolymerization inhibits directed VIF movement and causes the network to retract toward the nucleus,^28,29^ and inhibition or loss of kinesin-1 or cytoplasmic dynein alters VIF transport and distribution.^10–12,30^ Because kinesin-1 and dynein also move many membrane-bound organelles,^15^ a VIF defect after motor inhibition cannot, by itself, distinguish VIFs engaging motors directly from VIFs being displaced through association with another motor-driven structure. Indeed, when kinesin was first implicated in VIF organization, Gyoeva and Gelfand^10^ considered both possibilities, including that kinesin-driven membranous organelles might move the intermediate filaments anchored to them. To our knowledge, this organelle-mediated route has not previously been tested directly.

VIFs are increasingly viewed as a dynamic cytoplasmic network that interacts with the other cytoskeletal systems and helps position organelles.^31–34^ VIFs associate closely with mitochondria^18,35^ and form cage-like structures around lipid droplets^36,37^ and melanosomes,^17^ and in melanophores the VIF network disperses together with melanosomes. Whether organelles themselves move the filaments, however, has not been addressed. Such movement would be a form of hitchhiking, in which a cargo moves by associating with another cargo that has already recruited motors,^13^ as established for peroxisomes, lipid droplets, the ER, and RNA granules.^38–41^ Our results support organelle-associated transport of a subset of VIFs.

Two observations argue that VIF–mitochondria association does not simply reflect both structures being attached to a shared microtubule. Super-resolution imaging revealed VIFs crossing mitochondria without a detectable microtubule, and volume EM showed individual VIFs curving along mitochondrial surfaces at sites that only partly overlapped with mitochondria–microtubule contacts. The comigration we observed in living cells illustrates that the two structures can move together, but the causal evidence comes from the perturbation experiments.

How VIFs are physically coupled to mitochondria remains to be defined, but previous work points to two non-exclusive linkages. The first is direct: a region of the non-α-helical vimentin head domain localizes to mitochondria, and assembly-competent vimentin mutants lacking this region no longer restrain mitochondrial motility;^5^ recombinant vimentin also binds isolated mitochondria through its N-terminal region.^42^ The mitochondrial partner of this region is unknown, and because cryo-electron tomography places the head domains largely within the filament lumen,^43^ how it is accessed in assembled filaments remains an open question. The second linkage is bridged by the cytolinker plectin 1b, which is anchored in the outer mitochondrial membrane and links mitochondria to VIFs.^19^ The head-domain interaction is also regulated: phosphorylation of vimentin Ser-55 by the Rac1–PAK1 pathway releases mitochondria,^20^ providing a means by which only a subset of VIFs could be coupled at any given time. Disrupting either linkage should therefore reduce the VIF redistribution that accompanies mitochondrial repositioning.

Our perturbation experiments show that this association is functional. Depleting TRAK1, which is recruited to mitochondria by MIRO1 and links them to kinesin-1,^21,23^ shifted VIFs toward the nucleus together with mitochondria while leaving microtubule tracks intact, providing evidence that VIF distribution depends specifically on mitochondrial transport. Loss of function alone, however, cannot show that moving mitochondria is sufficient to move VIFs. Redirecting mitochondrial transport provided this evidence. Tethering a truncated kinesin-1 (residues 1–560) motor to mitochondria in cells lacking endogenous kinesin-1 drove mitochondria toward the cell periphery, and VIFs followed. Because this motor lacks the tail required for kinesin-1-dependent intermediate filament transport and acted only on mitochondria, the VIF redistribution indicates mediation by mitochondria. Consistent with this, mitochondria reached the periphery before VIFs accumulated at the same sites. Driving mitochondria toward microtubule minus ends had the opposite effect, carrying a substantial fraction of the VIF network toward the nucleus. Pasolli et al.^44^ addressed the converse question by moving VIFs directly and found that mitochondria only partly followed, suggesting that the coupling may be asymmetric, with motor-engaged mitochondria acting as vehicles for VIFs more readily than VIFs reposition mitochondria.

VIFs are generally regarded as anchors that restrain mitochondria: loss of VIFs increases mitochondrial motility, whereas VIF aggregation traps mitochondria,^5,26,45^ and vimentin similarly retains the ER and endolysosomes in the perinuclear region.^7^ Our results show that the relationship also runs in reverse: mitochondria driven by motors carry VIFs with them. These roles need not conflict. A single filament attached to a mitochondrion can be displaced when motors pull the organelle, whereas the interconnected network as a whole resists organelle movement; which effect dominates may depend on the balance between motor force and network resistance. Other intermediate filaments are also linked to mitochondria: loss of desmin or keratins mispositions mitochondria,^46,47^ and neurofilaments bind mitochondria directly.^48^ Moreover, in giant axonal neuropathy and in Charcot-Marie-Tooth disease caused by neurofilament light chain mutations, defective intermediate filament transport is accompanied by abnormal mitochondrial transport and distribution, attributed at least in part to trapping of mitochondria within filament aggregates. ^26,45,49^

Mitochondrial coupling does not, however, account for all VIF movement. VIF redistribution did not scale with mitochondrial redistribution: loss of kinesin-1 affected VIFs more than its effect on mitochondria would predict, and when mitochondria were driven toward the nucleus, part of the peripheral VIF network remained. This suggests that kinesin-1 also positions VIFs through routes that do not involve mitochondria, either by engaging VIFs directly or through an adaptor, ^12^ or by carrying them on other organelles such as the ER, which is also coupled to VIFs. ^7,44^ We therefore propose a hybrid model in which VIFs use multiple, non-exclusive modes of microtubule-dependent transport. ^32^ Such a mechanism would reconcile the strong dependence of VIF motility on microtubule motors with the scarcity of close VIF–microtubule contacts and would explain how individual VIFs within the same local network follow distinct trajectories,^9^ depending on which transport machinery each filament engages.

In summary, mitochondrial motility contributes to VIF transport and spatial organization, acting alongside direct or adaptor-mediated engagement of microtubule motors. These findings extend hitchhiking to a cytoskeletal filament and suggest that organelles can act as mobile intermediates through which motors shape the organization of the cytoskeleton.

## Materials and methods

### Plasmids

Rapalog-based (FRB–FKBP) chemical dimerization constructs ^25,50,51^ were used to acutely reposition mitochondria toward either the plus or minus ends of microtubules. For plus-end-directed transport, the motor domain of human kinesin-1 (KIF5B, residues 1–560) was fused to FRB and EGFP (KIF5B(1–560)–FRB–EGFP) in a pLVX backbone. For minus-end-directed transport, the plant kinesin-14 (from *Physcomitrella patens*)^27^ motor construct FRB–mStayGold–Kinesin14VIb was cloned into a pTwist backbone. To permit rapalog-dependent recruitment of either motor to mitochondria, FKBP and mCherry were fused to the C-terminal 31-amino-acid mitochondrial outer-membrane-targeting sequence of human mitochondrial antiviral-signaling protein (MAVS; residues 510–540),^52^ generating FKBP–mCherry–OMM in a pLVX backbone. These constructs were commercially synthesized by Twist Bioscience or Synbio Technologies.TRAK1 knockdown was achieved using MISSION lentiviral shRNA constructs purchased from Millipore Sigma (TRCN0000036275 and TRCN0000421920). Both constructs contain a puromycin-resistance cassette.

### Antibodies

Primary antibodies: chicken anti-vimentin (BioLegend, #919101 and EnCor Biotechnology, #CPCA-Vim), mouse anti-α-tubulin (clone DM1A, produced in-house and Thermo Fisher Scientific, #62204), rabbit anti-TOM20 (Proteintech, #11802-1-AP and Abcam, #ab78547), and rabbit anti-TRAK1 (Thermo Fisher Scientific, #PA5-44180). Secondary antibodies: FITC-conjugated goat anti-mouse IgG (#115-095-003), Rhodamine Red-X (TRITC)-conjugated AffiniPure goat anti-rabbit IgG (H+L) (#111-025-144), and Alexa Fluor 647-conjugated donkey anti-chicken IgY (#703-605-155) from Jackson ImmunoResearch. Goat anti-chicken Alexa Fluor Plus 488 (A32931), goat anti-mouse Alexa Fluor Plus 555 (A32727; both Thermo Fisher Scientific) and an Alexa Fluor 405 anti-rabbit secondary antibody.

### Cell culture and transfection

Human retinal pigment epithelial (RPE) cells, human BJ05 fibroblasts, and African green monkey COS-7 cells were cultured in DMEM (Sigma-Aldrich, #D5648) supplemented with 10% fetal bovine serum. Cells were maintained at 37 °C in a humidified atmosphere containing 5% CO₂. TRAK1 knockdown was achieved by transient or stable expression of the indicated shRNA constructs. For stable knockdown, lentivirus was produced in HEK293FT cells, and transduced cells were selected with puromycin. Single-cell clones were isolated by limiting dilution. Knockdown efficiency was validated by measuring TRAK1 mRNA levels using qPCR, and the expected mitochondrial aggregation phenotype was confirmed by immunofluorescence microscopy. The kinesin-1-knockout RPE cell line was described previously.^12^ For imaging experiments, cells were seeded onto cleaned glass coverslips in 35-mm culture dishes. The following day, cells were transfected with plasmid DNA using ViaFect Transfection Reagent (Promega, #E4981) according to the manufacturer’s instructions. For transient TRAK1-knockdown experiments, cells were fixed 48 h after transfection. For all other experiments, cells were fixed or imaged 24 h after transfection. For chemically induced dimerization experiments, cells were treated with 100 nM rapalog heterodimerizer A/C (TakaraBio, #635057) 24 h after transfection. Cells were incubated with heterodimerizer for 2 h, then cells were washed and fixed for immunostaining.

### Live cell spinning disk confocal microscopy

Live mEmerald-vimentin/mTagRFP-T-tubulin knock-in RPE cells were labeled with MitoTracker Deep Red and imaged on a Nikon Ti spinning-disk confocal microscope using an SR Apo TIRF 100×/1.49 NA oil objective and an Andor DU-897 EMCCD camera. Acquisition was controlled by NIS-Elements AR 4.40.00. Vimentin and mitochondria were excited at 488 and 640 nm, with exposures of 70 and 10 ms, respectively. Images were acquired at approximately 3 s intervals with a pixel size of 0.134 µm. Images were denoised using the AI denoising tool in Nikon NIS-Elements 5.42.07. A fixed background was estimated for each channel from the temporal median using a minimum filter and Gaussian smoothing (σ = 3 µm); 60% of this background was subtracted from every frame. Display limits and gamma were held constant over time (1.0 for mitochondria and 0.75 for vimentin). Leading and trailing mitochondrial ends were identified in the mitochondrial channel and marked with vector arrows at corresponding positions in the merged and separate-channel panels. Movies retain all 13 acquired time points over 36 s and use the recorded timestamps at fourfold playback speed.

### Fixed cell STED microscopy

COS-7 cells were fixed with 4% paraformaldehyde and 0.2% glutaraldehyde for 15 min, washed three times in PBS, and treated with 0.1% sodium borohydride in PBS for 7 min. After washing, cells were permeabilized and blocked in 1% BSA containing 0.25% Triton X-100. The same BSA/Triton solution was used for antibody staining. Images were acquired on a Leica SP8 STED microscope equipped with an HC PL APO CS2 100×/1.40 NA oil-immersion objective. Vimentin and α-tubulin were excited at 499 and 552 nm, with 592 and 775 nm depletion, respectively, and gated HyD detection (0.5–6.0 ns). TOM20 was excited at 405 nm and acquired in confocal mode. Single optical sections were acquired at approximately 12.6 nm pixel sampling and a 700 Hz scan rate.

Images were processed using custom Python scripts. A slowly varying background was estimated for the filament channels using a 1.5 µm minimum filter followed by Gaussian smoothing (σ = 3 µm), subtracted, and negative values set to zero. Gaussian smoothing of 30 nm was then applied to the filament images. TOM20 images were smoothed with a 45 nm Gaussian. Filament-shaped crosstalk in the TOM20 channel was corrected by subtracting shifted, blurred filament templates fitted in mitochondria-free cytoplasm. Channel-specific intensity limits were fixed for each field; TOM20 was displayed linearly, and gamma values of 0.75 and 0.75–0.90 were used for vimentin and microtubules, respectively. Three-color composites used a half-weight microtubule channel and intensity-dependent blending of vimentin over the combined mitochondrial and microtubule signals. Individual channels and pairwise merges were retained for the same regions.

### Fixed cell spinning disk confocal microscopy

Immunofluorescence staining was performed as described previously.^9^ Cells were pre-fixed and simultaneously permeabilized for 3 min at 37 °C in PHEM buffer containing 0.3% glutaraldehyde and 0.2% Triton X-100. PHEM buffer consisted of 60 mM PIPES, 27 mM HEPES, 10 mM EGTA, and 8 mM MgSO₄·7H₂O, adjusted to pH 7.0. Cells were subsequently fixed for 15 min at 37 °C in 4% paraformaldehyde prepared in PHEM buffer. Residual aldehyde groups were quenched with sodium borohydride (5 mg/ml) for 5 min, and cells were washed three times for 5 min each in PBS, then blocked in blocking buffer (2.5% BSA with 0.1% Triton X-100). Cells were incubated with primary and secondary antibodies diluted in blocking buffer for 45 min each at room temperature. Coverslips were mounted and cured for 24 h at room temperature before imaging. Images were acquired using spinning-disk confocal systems, a Nikon Eclipse Ti2 stand equipped with a Yokogawa CSU-W1 spinning-disk confocal head (50-μm pinhole) and a Hamamatsu ORCA-Fusion digital CMOS camera (C13440-20CU), controlled by Nikon NIS-Elements software (version 5.42.01). System was fitted with a 60×/1.49 NA or 100×/1.45 NA oil-immersion objective. Z-stacks were acquired at 0.2-μm intervals, and raw images were used directly for downstream image analysis.

### Image processing and analysis

Fluorescence Z-stacks were processed using a custom Python pipeline (Python 3.10, NumPy, SciPy, pandas, scikit-image, and Matplotlib). Nuclei were segmented by three-dimensional Gaussian smoothing and Otsu thresholding, and the cytoplasmic mask was defined as a manually traced cell boundary, extruded through the Z-stack and excluding the nuclear mask. For each cytoplasmic voxel, a normalized radial coordinate r̃ (0 = nuclear boundary, 1 = cell periphery) was calculated from its relative distance to the nucleus and cell edge, and background-subtracted channel intensities were binned along r̃ to generate per-cell radial distribution profiles, summarized by the intensity-weighted radial center of mass (rCOM), radial standard deviation (sigma), and perinuclear, intermediate, and peripheral signal fractions. Condensation/compactness was quantified as the effective intensity volume (background-corrected signal normalized to cytoplasmic volume) and the Gini coefficient of the intensity distribution. All segmentations and radial maps were visually quality controlled per cell, and cells failing predefined exclusion criteria (segmentation failure, incomplete axial sampling, saturation) were excluded before analysis. Full analysis code and parameter definitions are available at request. Generative artificial intelligence (OpenAI Codex) was used to assist with the development, debugging, and documentation of the custom Python image-analysis code.

### FIB-SEM reconstruction and proximity analysis

We reanalyzed the perinuclear COS-7 FIB-SEM volume described by Renganathan et al.^6^. Sample preparation, acquisition at 2 nm isotropic sampling and manual filament tracing were performed as described in that study. Filament paths were checked against the original tracing graph, corrected where necessary, and resampled and smoothed using the published procedure. VIFs and microtubules were modeled as tubes of 6 and 12 nm radius, respectively. Mitochondrial surfaces were reconstructed from the existing 8 nm isotropic segmentation mask.

Mitochondrial surface proximity was calculated from the shortest distance between native surface-triangle centroids and filament centerline segments, minus the filament radius. Negative gaps were set to zero, and cumulative distributions were weighted by triangle area. Filament proximity was calculated at centerline-segment midpoints relative to the nearest native mitochondrial surface, weighted by segment length. Distances were signed according to whether the midpoint lay inside or outside the mitochondrial mask before subtracting filament radius. For cumulative filament-length distributions, negative gaps were excluded from the numerator but retained in the total-length denominator. Surface centroids were restricted to at least 100 nm from crop boundaries; filament measurements required both the midpoint and nearest mitochondrial surface point to meet this criterion. Artificial closure surfaces were excluded from measurements. All proximity measurements derive from one reconstructed cell volume.

### Three-dimensional rendering

Reconstructions were rendered in Blender 4.5 using Cycles and orthographic projection. Mitochondrial components were smoothed independently (σ = 16 nm) for display; measurements used the unsmoothed native surface. Reciprocal distance maps were displayed from 0 to 200 nm, with larger gaps shown in white and modeled overlaps assigned to zero. The selected mitochondrion and associated VIF were shown using their complete in-volume geometry. Surrounding structures were faded for context, and the segmented basal plasma membrane provided an anatomical reference.

### Statistical analysis

For each analysis, the number of biological replicates and the total number of cells analyzed are reported in the corresponding results text. Statistical significance was assessed using the Mann-Whitney U test. Significance is denoted as ns (not significant, p ≥ 0.05), * p < 0.05, ** p < 0.01, *** p < 0.001, **** p < 0.0001.

## Supporting information

Supplementary movie 1, mitochondria and Vimentin IF motility in live cell

Supplementary movie 2, mitochondria and Vimentin IF moving reciprocal

Supplementary movie 3, mitochondria and microtubule moving reciprocal

## Statements and Declarations

All authors declare no conflict of interest.

## Acknowledgements and Funding Information

The Gelfand lab is supported by the grant R35- GM131752 from National Institute of General Medical Sciences. The Jennifer Lippincott-Schwartz lab is supported by the Howard Hughes Medical Institute Janelia Research Campus. We thank the CellMap Project Team at HHMI’s Janelia Research Campus for generating and sharing the FIB-SEM dataset used in this study.

## Data Availability Statement

All data and analysis code reported in this paper will be shared upon request.

## Supplementary figure legends

**Figure S1.**
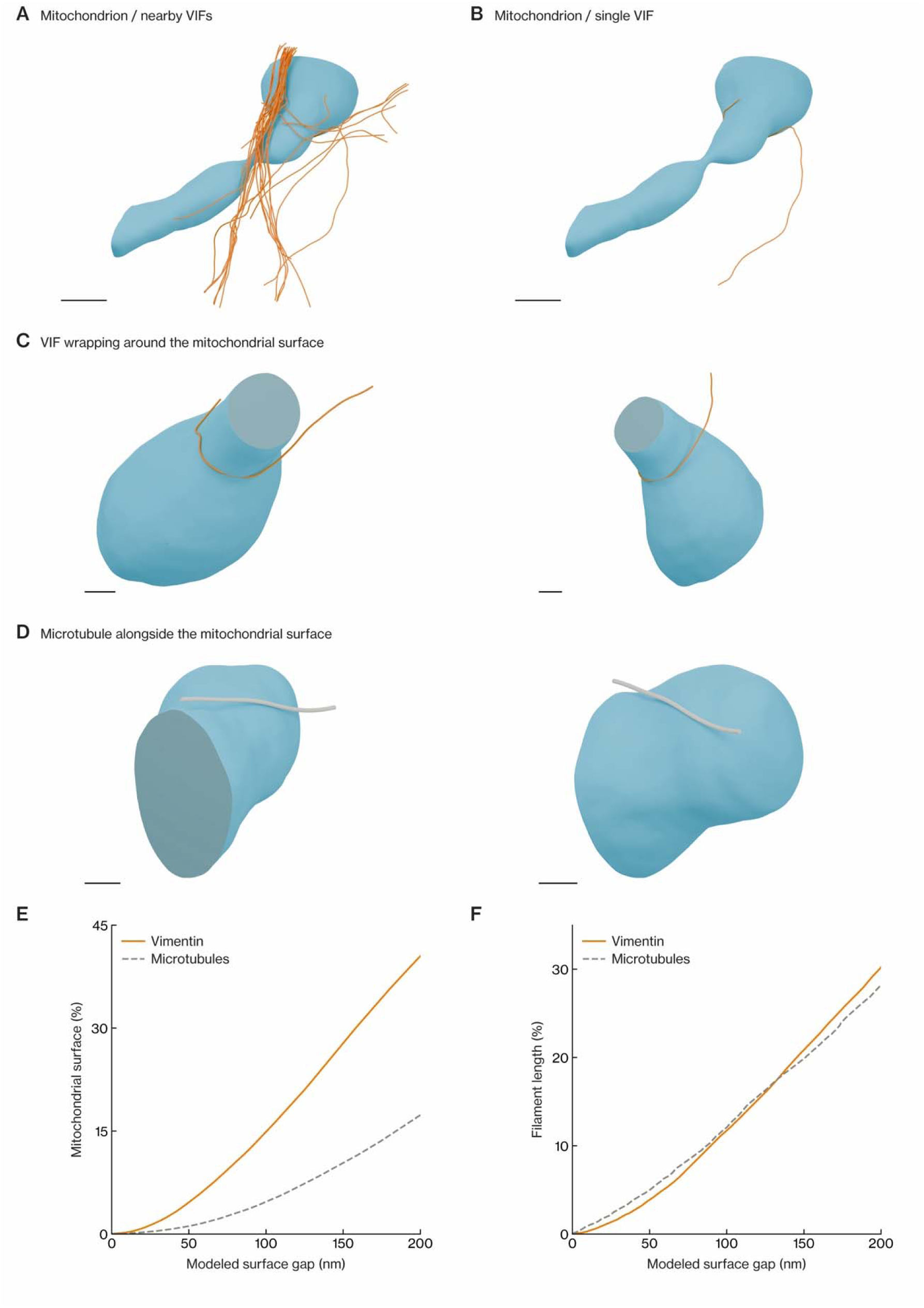
VIFs and microtubules follow the mitochondrial surface. (A) A segmented mitochondrion (light blue) with nearby VIFs (orange). (B) The same mitochondrion with a single VIF, viewed from the same direction. (C) Two additional views of the region where this VIF curves around the mitochondrial surface. (D) Two views of a separate mitochondrion with an adjacent microtubule (gray). Other structures are omitted. Clipping planes expose the regions in C and D; flat surfaces mark display cuts or boundaries of the imaged volume. (E and F) Cumulative mitochondrial surface area (E) and filament length (F) within the indicated modeled gap of the other structure. Measurements exclude a 100-nm volume margin and use modeled filament radii of 6 nm for VIFs and 12 nm for microtubules. In E, modeled overlap counts at zero; in F, negative gaps are excluded from the cumulative length but retained in the total length. All panels derive from one COS-7 cell volume. Scale bars: 500 nm (A and B); 200 nm (C and D).

**Figure S2.**
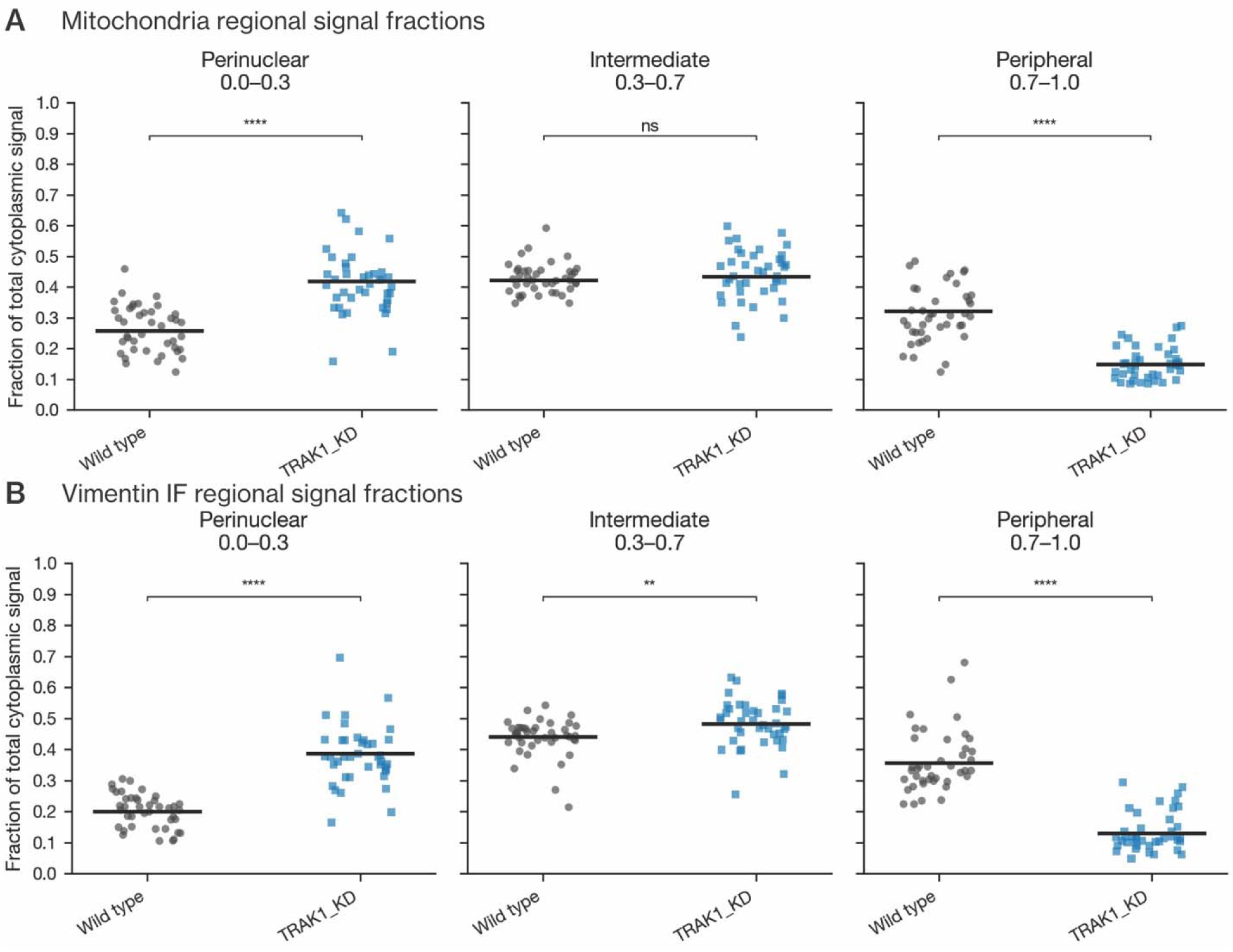
Regional signal fractions in TRAK1 KD RPE cells. (A and B) Perinuclear (0–0.3), intermediate (0.3–0.7), and peripheral (0.7–1.0) signal fractions of mitochondria (A) and VIFs (B) in wild-type and TRAK1 KD RPE cells. Each point is one cell; bars show the mean. N = 41 wild-type and 40 TRAK1 KD cells from three independent experiments.

**Figure S3.**
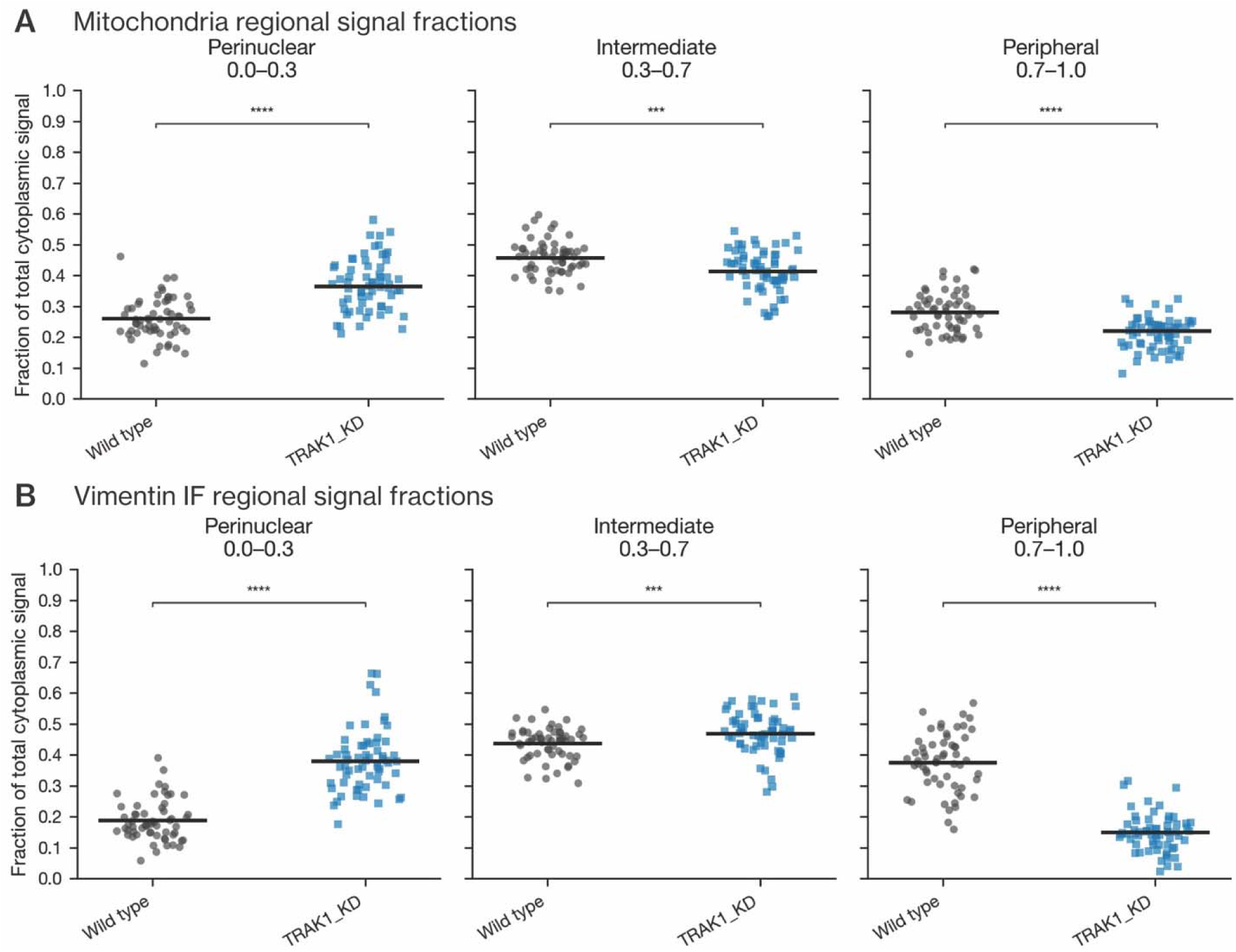
Regional signal fractions in TRAK1 KD BJ05 fibroblasts. (A and B) Perinuclear (0–0.3), intermediate (0.3–0.7), and peripheral (0.7–1.0) signal fractions of mitochondria (A) and VIFs (B) in wild-type and TRAK1 KD BJ05 cells. Each point is one cell; bars show the mean. N = 58 wild-type and 63 TRAK1 KD cells from three independent experiments.

**Figure S4.**
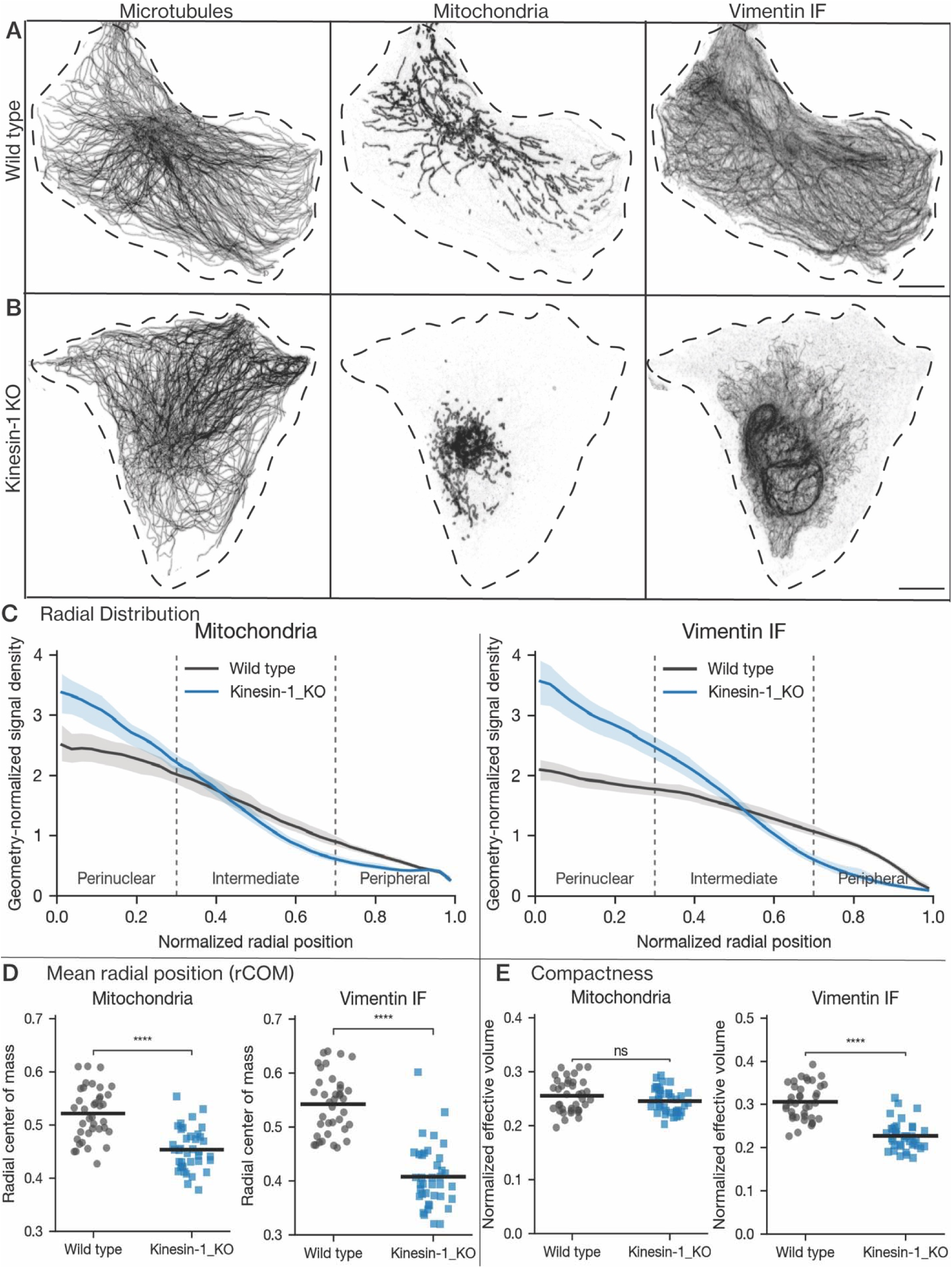
Loss of kinesin-1 shifts mitochondria and VIFs toward the perinuclear region. (A and B) Representative inverted-contrast immunofluorescence images of microtubules (left), mitochondria (middle), and VIFs (right) in wild-type (A) and kinesin-1 KO (B) RPE cells. Dashed lines outline the cell boundary. (C) Geometry-normalized radial profiles of mitochondria (left) and VIFs (right) in wild-type (black) and kinesin-1 KO (blue) cells. (D) Mean radial position (rCOM) of mitochondria (left) and VIFs (right). (E) Compactness (normalized effective intensity volume) of mitochondria (left) and VIFs (right). Each point is one cell; bars show the mean. Regional signal fractions are shown in Fig. S5. Scale bar, 10 µm. N = 38 wild-type and 38 kinesin-1 KO cells from three independent experiments.

**Figure S5.**
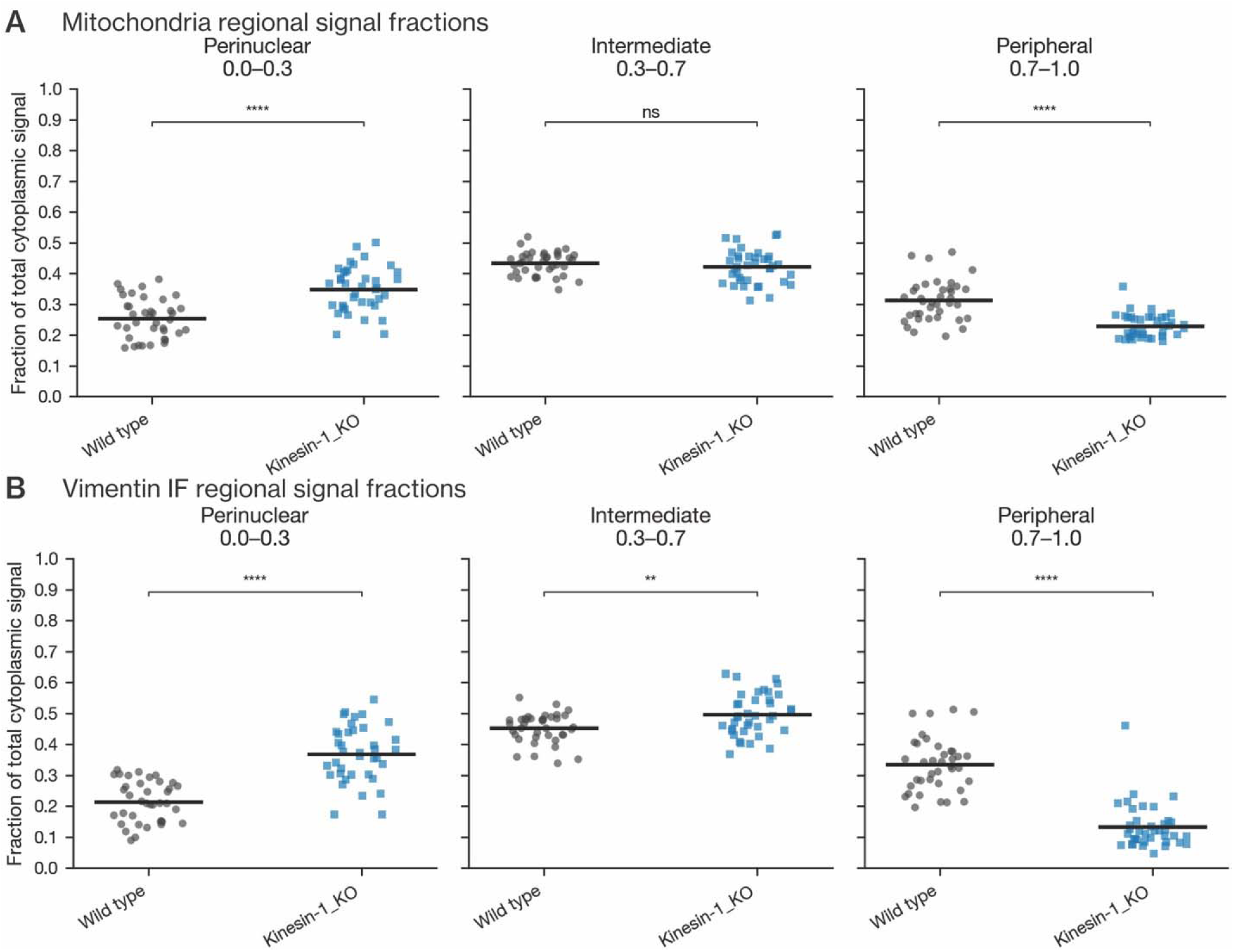
Regional signal fractions in kinesin-1 KO RPE cells. (A and B) Perinuclear (0–0.3), intermediate (0.3–0.7), and peripheral (0.7–1.0) signal fractions of mitochondria (A) and VIFs (B) in wild-type and kinesin-1 KO cells. Each point is one cell; bars show the mean. N = 38 wild-type and 38 kinesin-1 KO cells from three independent experiments.

**Figure S6.**
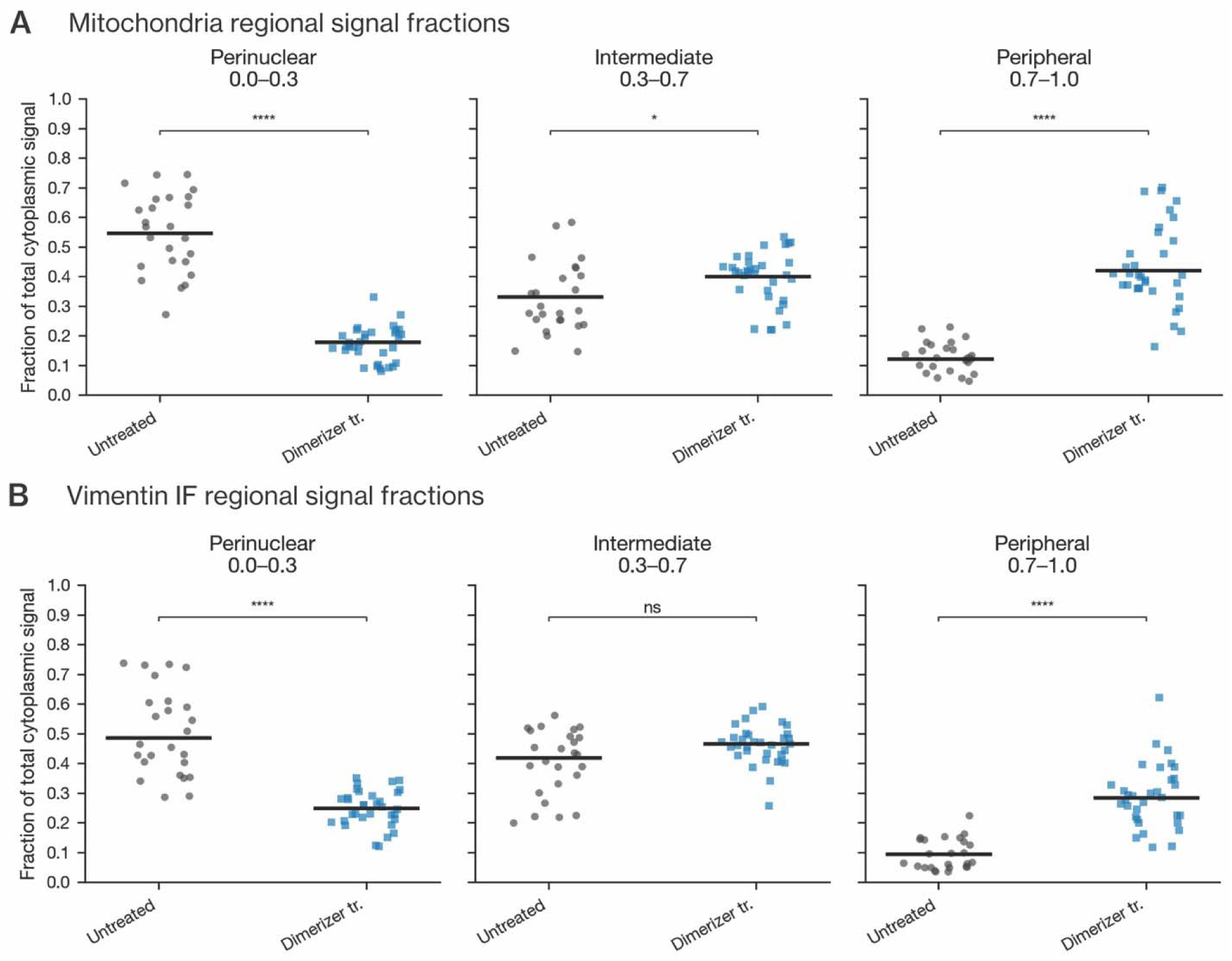
Regional signal fractions after recruitment of truncated kinesin-1 to mitochondria. (A and B) Perinuclear (0–0.3), intermediate (0.3–0.7), and peripheral (0.7–1.0) signal fractions of mitochondria (A) and VIFs (B) in untreated and dimerizer-treated kinesin-1 KO RPE cells expressing truncated kinesin-1–FRB–GFP and OMM–FKBP–mCherry. Each point is one cell; bars show the mean. Images, radial profiles, mean radial position, and compactness are shown in Fig. 6. N = 25 untreated and 33 dimerizer-treated cells from three independent experiments.

**Figure S7.**
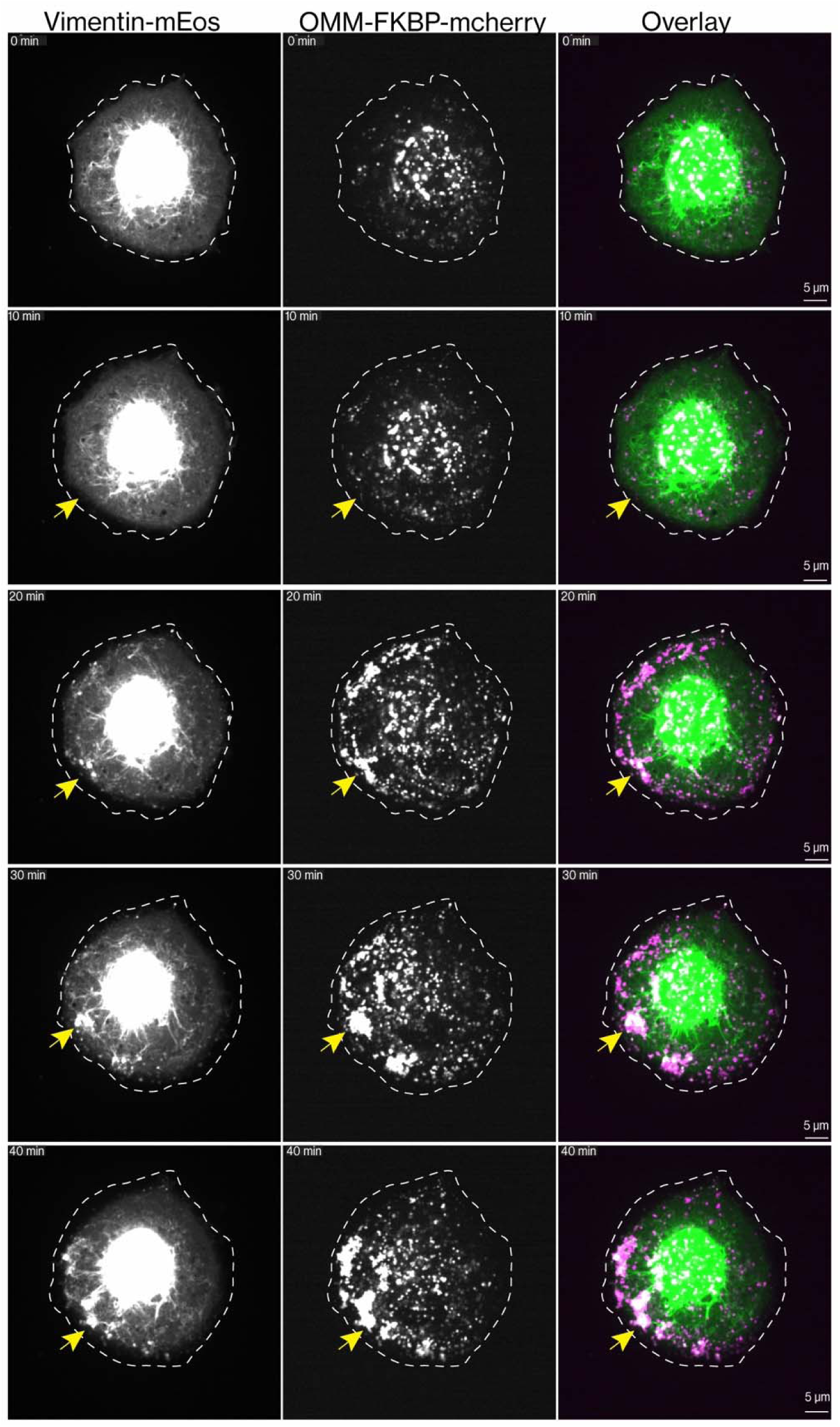
Live-cell snapshots of VIFs and mitochondria after recruiting truncated kinesin-1 to mitochondria in Kinesin-1 -knockout cells. Kinesin-1-knockout RPE cell expressing vimentin–mEos, OMM–FKBP–mCherry and truncated kinesin-1–FRB. Vimentin–mEos was imaged in its native green (non-photoconverted) state. Before dimerizer addition (0 min), VIFs and mitochondria were concentrated around the nucleus. Adding 100 nM dimerizer induces FKBP–FRB heterodimerization, which recruits the plus-end-directed kinesin-1 motor domain to the mitochondrial outer membrane. Times are relative to dimerizer addition; the 0-min image was acquired immediately before addition. Mitochondria moved toward the cell periphery and formed peripheral clusters, visible by 20 min. Vimentin then accumulated at the same sites and was prominent by 30 min. This indicates that mitochondrial redistribution precedes the redistribution of VIFs. Yellow arrows mark the same peripheral region over time. In the overlays, vimentin–mEos is green and mitochondria are magenta. The dashed line marks the cell outline at 0 min and is overlaid unchanged on later time points. Display contrast was set to show peripheral filaments, so the perinuclear vimentin signal is saturated. Images are single confocal planes. Scale bars, 5 µm.

**Figure S8.**
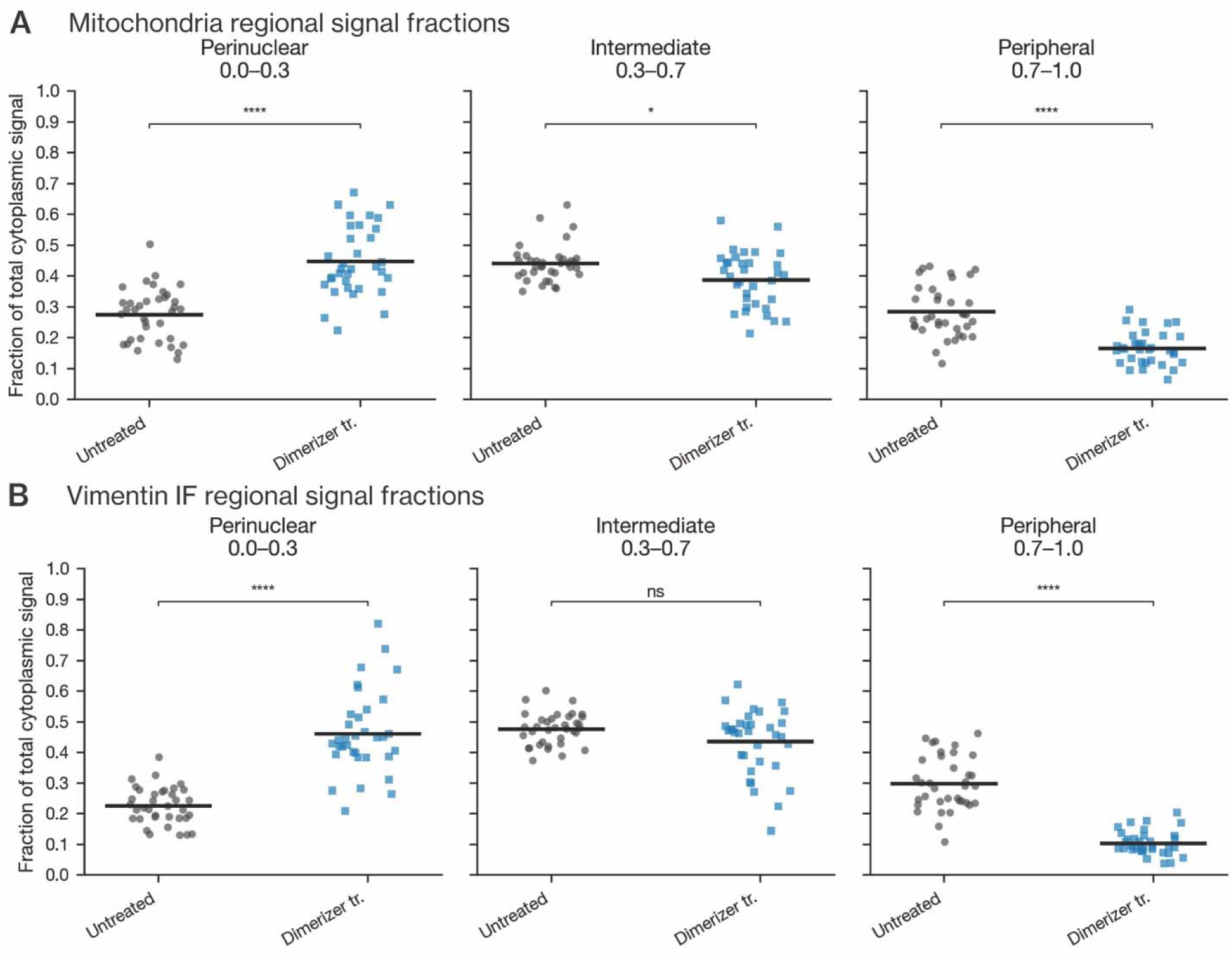
Regional signal fractions after recruitment of kinesin-14 VIb to mitochondria. (A and B) Perinuclear (0–0.3), intermediate (0.3–0.7), and peripheral (0.7–1.0) signal fractions of mitochondria (A) and VIFs (B) in untreated and dimerizer-treated wild-type RPE cells expressing kinesin-14 VIb–FRB–GFP and OMM–FKBP–mCherry. Each point is one cell; bars show the mean. Images, radial profiles, mean radial position, and compactness are shown in Fig. 7. N = 36 untreated and 34 dimerizer-treated cells from three independent experiments.

**Figure S9.**
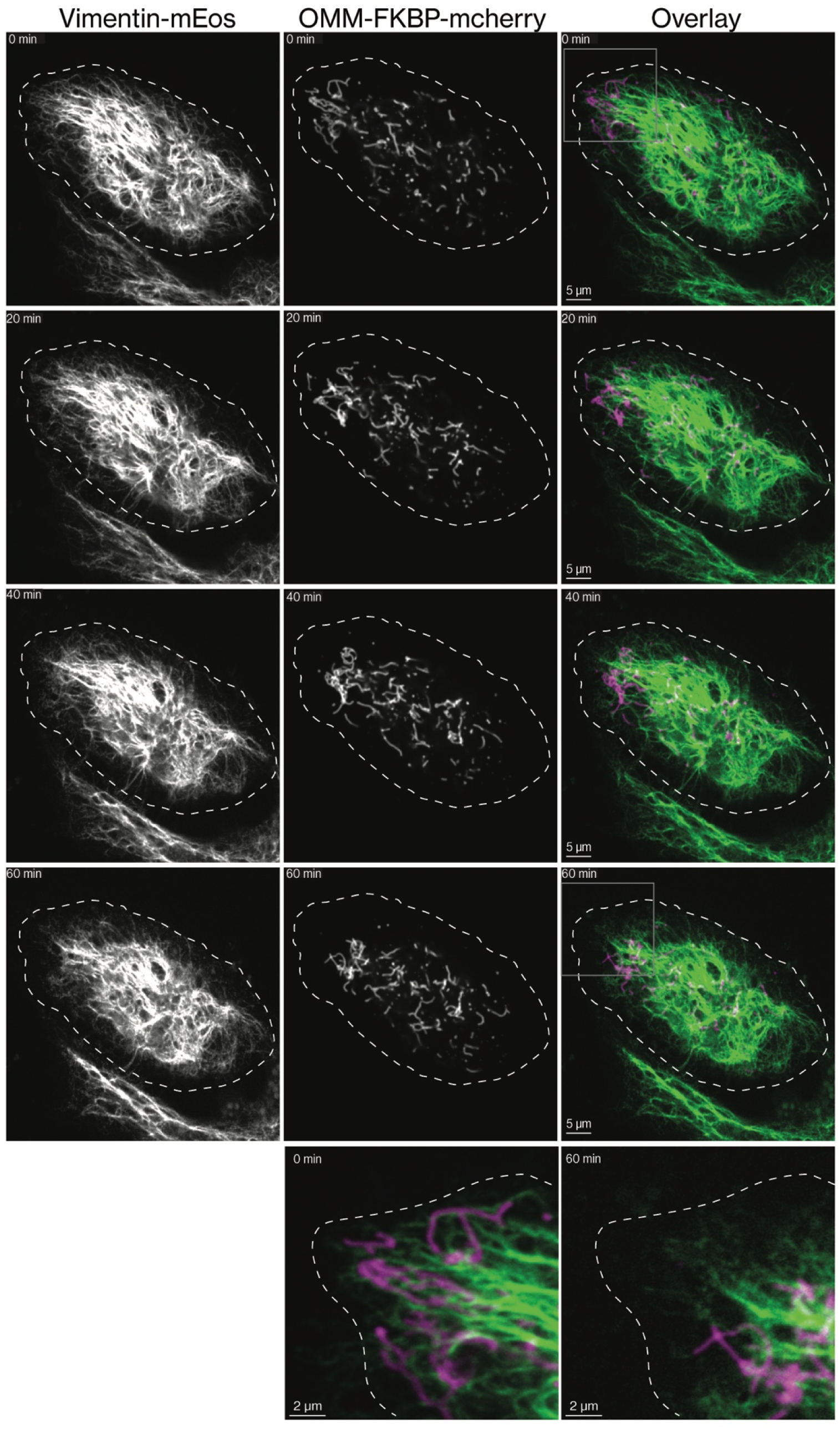
Live-cell snapshots of VIFs and mitochondria after recruiting kinesin-14 VIb to mitochondria. RPE cell expressing vimentin–mEos, OMM–FKBP–mCherry and kinesin-14 VIb–FRB. Vimentin–mEos was imaged in its native green (non-photoconverted) state. Adding 100 nM A/C heterodimerizer induces FKBP–FRB heterodimerization, which recruits the minus-end-directed motor kinesin-14 VIb to the mitochondrial outer membrane. Times are relative to dimerizer addition; the 0-min image was acquired immediately before addition. Over 60 min, mitochondria moved progressively toward the nucleus, clearing the peripheral cytoplasm, most evidently in the gray boxed region. Over the same period, the peripheral vimentin network thinned and retracted from the cell edge, leaving a more compact perinuclear network. In the adjacent cell at the bottom of the field, which expresses vimentin–mEos but not the recruitment constructs, the vimentin network is unchanged over the same period. This argues against photobleaching as the cause of the peripheral vimentin loss. In the overlays, vimentin–mEos is green and mitochondria are magenta. The dashed line marks the cell outline at 0 min and is overlaid unchanged on later time points. Boxed regions at 0 and 60 min are enlarged at the bottom (overlay). Images are single confocal planes. Scale bars: 5 µm (whole cell), 2 µm (enlargements).

## Movie Captions

**Movie 1. Mitochondria-VIF co-motility**

Spinning-disk confocal time-lapse of a live RPE cell with endogenously tagged vimentin-mEmerald and labeled with MitoTracker Deep Red. The merge shows mitochondria in blue-cyan and vimentin in orange; individual channels are shown in grayscale Filled green and open magenta arrows mark the leading and trailing mitochondrial ends during the outward movement. The same positions are marked in all three panels, with end identities retained during the return movement. All 13 acquired time points are shown, spanning 36 s. Playback, 4 times acquisition speed. Scale bars, 2 µm.

**Movie 2. Volume EM reconstructions of mitochondria and VIFs featuring reciprocal Mito-VIF distance maps.**

The movie displays a 3D rendering of mitochondria (light blue), VIFs (orange) and microtubules (light gray) as depicted in the main figure. The camera pivots from the head-on view at the outset to the top-down view by the midpoint before returning to the original head on view by the final frame. As the movie begins, the microtubule mask is hidden to better display the mitochondria-VIF interactions. At approximately 00:10, the orange VIFs fade from view, and a mitochondria-to-VIF distance map is displayed on the surface of all mitochondria (orange to white, legend displayed on movie). At 00:20 the original mitochondria and VIF masks return revealing the two structures from the top-down perspective. As the movie continues, the camera pivots back down to the head-on view, and at approximately 00:25 the mitochondria fade to faint, low transparency traces and the VIF-to-mitochondria distance map is projected onto the surface of individual VIFs (blue to white, legend displayed on movie). The basal plasma membrane provides anatomical context. The distance key is shown only when each map is fully displayed Scale bar, 500 nm.

**Movie 3. Volume EM reconstruction of mitochondria and microtubules featuring reciprocal Mito-MT distance maps**

Microtubules and mitochondrial surfaces in the same COS-7 FIB-SEM reconstruction. The movie follows the same camera path and display sequence, isolating mitochondria and microtubules after the three-component overview. The mitochondrial surface map shows the distance map to microtubules, using neutral gray at 0 nm and white at 200 nm and above. The reciprocal microtubule map shows the gap to the mitochondrial surface in light blue at 0 nm, fading to white at 200 nm and above, with mitochondria retained as faint neutral gray context. Structural microtubules are gray. Camera motion, color transitions and acquisition-boundary closures have the same interpretation as in the VIF movie. Scale bar, 500 nm.

